# Anti-PD-1-iRGD Peptide Conjugate Boosts Antitumor Efficacy via Engagement Augmentation and Penetration Enhancement of T cells

**DOI:** 10.1101/2023.08.04.551949

**Authors:** Yunfeng Pan, Qi Xue, Yi Yang, Tao Shi, Hanbing Wang, Xueru Song, Xueyi Yang, Baorui Liu, Zhentao Song, Jie P. Li, Jia Wei

## Abstract

Despite the important breakthroughs of immune-checkpoint inhibitors (ICIs) in recent years, the overall objective response rate (ORR) remains limited in various cancers. Here, we synthesized programmed cell death protein-1 (PD-1) antibody iRGD conjugate (αPD-1-(iRGD)_2_) through glycoengineering and bio-orthogonal reaction. αPD-1-(iRGD)_2_ exhibited extra iRGD receptor dependent affinity to several cancer cell lines rather than normal cell lines. Via dual targeting, αPD-1-(iRGD)_2_ engageed tumor cells and T cells thus mediating T cell activation and facilitating tumor elimination. Besides, the attachment of iRGD impressively improved the penetrability of both PD-1 antibody and PD-1^+^ T cells. In multiple syngeneic mouse models, αPD-1-(iRGD)_2_ effectively reduced tumor growth with satisfactory biosafety. Moreover, results of flow cytometry and single-cell RNA-seq revealed that αPD-1-(iRGD)_2_ remodeled the tumor microenvironment (TME) and expanded a unique population of “better effector” CD8^+^ tumor infiltrating T cells (TILs) expressing stem and memory associated genes including *Tcf7*, *Il7r*, *Lef1* and *Bach2*. Conclusively, αPD-1-(iRGD)_2_ could be a novel and promising therapeutic approach for cancer immunotherapy.

**Statement of significance:** Designed against the clinical dilemma of unsatisfied response rate after contemporary cancer immunotherapy, αPD-1-(iRGD)_2_ engages T cells and tumor cells, promotes T cell infiltration and expands a unique population of “better effectors” with enhanced therapeutic potential for the treatment of cancer.

## Introduction

Immunotherapy especially immune checkpoint inhibitor (ICI) brought significant survival benefit for advanced cancer patients(1-3). Although indications of ICIs have expanded over time, the ORR remains merely 10-30% among several solid tumors(4-6). Researchers endeavored to develop novel ICIs with better antitumor efficacy. BiTEs are emerging T cell-engaging therapies with promising cancer treatment potential(7). Due to their affinity to both T cells and tumor cells, BiTEs exhibit unique engagement effects and mediate effective tumor control at lower doses compared with antibodies(8). Traditionally, BiTEs contains one or more CD3 binding domain to realize T cell engaging, which may induce unselective T cell activation and severe side effects(9). Additionally, researchers found that PD-1^+^ T cells enriched more tumor specificity(10). Therefore, we attempted to design a new form of BiTE based on PD-1 antibodies. However, analogous to Chimeric Antigen Receptor T-Cell Immunotherapy (CAR-T), the success of BiTEs realized in hematologic neoplasms has yet to be obtained in solid tumors. The difficulty of penetration caused by dense microenvironment plays a role(8).

ICIs represented by PD-1/PD-L1 pathway inhibitors currently prescribed in clinic all belong to antibodies with a relative molecular mass of around 150 kDa(11). The huge scale impedes the distribution of ICIs in tumor bulk. Recently, a positron emission tomography (PET) imaging study with zirconium-89 (89Zr)-labeled pembrolizumab illustrated that antibodies required approximately 5-7 days to reach peak concentration in tumor region(12, 13). Meanwhile, multidimensional angiography demonstrated that antibodies distributed primarily in perivascular area after systemic administration, exclusively occupying around 35% of tumor volume(14). The antitumor efficacy of PD-1 monoclonal antibody (mAb) depends on sufficient infiltration of T cells (15). Slow and uneven local tumor distribution limits the efficacy of PD-1 monoclonal antibody. In a small sample scale clinical trial, the local maximum standardized uptake value (SUVmax) of PD-1 antibody was positively correlated with the patients’ response to immunotherapy, progression-free survival (PFS) and overall survival (OS)(12). However, there is no correlation between clinical response and PD-1/PD-L1 immunohistochemical (IHC) staining scores, suggesting reforming penetrability of PD-1 mAb is a potential strategy to improve the efficacy of ICIs. Hence, there is an urgent need to develop general, effective and safe antibody penetration enhancement strategy.

Internalizing RGD (iRGD, c(CRGDKGPDC)), a tumor penetrating peptide, has been applied to deliver agents, including chemotherapeutics, single-chain mAbs and nanoparticles, towards tumor bulk through interacting successively with integrins and Neuropilin-1 (NRP1)(16, 17). Theoretically, the attachment of iRGD not only provides αPD-1 with binding affinity to integrins and NRP1, but also facilitates tumor penetration. Our team has reported that T cell membrane modification with iRGD impressively strengthened T cell penetration and antitumor efficacy(18, 19). However, due to technique restriction, iRGD modification of native double-chain mAb hasn’t been reported. Compared with single chain variable fragment (scFv) and other fragments of mAb, complete natural antibody possesses a longer half-life and profound bioactivities(20). Traditionally, iRGD modification was conducted through plasmid transfection and subsequent eukaryotic or prokaryotic expression. The prokaryotic system cannot express double-chain mAb, while the eukaryotic system has difficulty in constructing the cyclic structure of iRGD(21, 22). Hence, we turn to nongenetic glycoengineering of the conserved glycosylation site of the Fc domain. In our previous work, reducing the complexity of Fc glycan could dramatically accelerate the reaction. Therefore, we established LacNAc-conju platform, a one-step site-specific native IgG modification technique(23). Based on a native mAb, LacNAc-conju platform allows for site-specific attachment of iRGD peptide with a drug antibody ratio (DAR) close to 2.

In this study, we generated αPD-1-(iRGD)_2_ through glycoengineering and bio-orthogonal reaction. iRGD modification largely refined the penetration of both PD-1 mAb and T cells. Besides, similar to BiTEs, αPD-1-(iRGD)_2_ engages T cells and tumor cells thus promoting tumor elimination. Systemic administration of αPD-1-(iRGD)_2_ elicited TME remodulation and impressive tumor growth control in various mouse tumor models. Moreover, αPD-1-(iRGD)_2_ activates and expands a unique subset of TILs, expressing stem and memory associated genes (*Tcf7*, *Il7r*, *Lef1* and *Bach2*), which may be the potential mechanism of impressive tumor regression induced by αPD-1-(iRGD)_2_.

## Results

### Synthesis and characterization of αPD-1-(iRGD)_2_

An anti-PD-1 antibody, CS1003, was selected for its cross-affinity to both human and mouse PD-1. Although the whole synthesis procedure of αPD-1-(iRGD)_2_ was accomplished in a one-step reaction, we would split the reaction into several parts and describe them separately in a logical order (**Figure 1a**). Firstly, the glycan structure on the N297 residue of the antibody CS1003 was modified by removing redundant sugars using EndoS (an endoglycosidase for specific hydrolysis at N297) and trimming α-(1,6)-fucose with AlfC (an α-fucosidase). Then, bovine β-1,4-galactosyltransferase 1 (Y289L) (β4GalT1) was used to add a galactose molecule to the GlcNAc residue, creating a monoantennary disaccharide structure, LacNAc, on N297 of CS1003. To meet the one-step procedure of this platform, we introduced the GDP-fucose group to the N_3_-iRGD peptide via click chemistry (**Figure 1b** and Figure S1a). Electro Spray Ionization-Mass Spectroscopy (ESI-MS) manifested the relative molecular mass of core substrate GDP-FAmP4Prop (845.1989 Da), which was within the allowable range of error (Figure S1b). The donor substrate of FT enzyme, GDP-Fuc-iRGD, was purified through Prep-HPLC and ESI-MS indicated its relative molecular mass as 936.7961 Da (Figure S1c). Then, using the FT enzyme and GDP-Fuc-iRGD, we nearly fucosylated all of the LacNAc units on CS1003 (≥1.8 iRGD per antibody) under saturation conditions (**Figure 1c**). ESI-MS confirmed the relative molecular weight of αPD-1-(iRGD)_2_ was 148833.00 Da. For the convenience of subsequent *in vivo* distribution assay and fluorescence imaging, we attached extra fluorophores, Cy5, to αPD-1-(iRGD)_2_ (αPD-1-(iRGD)_2_-Cy5) and αPD-1 (αPD-1-Cy5). After synthesis procedure, characterization assay was conducted to ensure the stability and consistence of PD-1 mAb. Generally, ELISA showed no difference in binding affinity towards both human and murine PD-1 protein between αPD-1-(iRGD)_2_ and the unmodified αPD-1 (**Figure 1d**). Calculated dissociation constant (Kd) of αPD-1-(iRGD)_2_ to human PD-1 was 2.855, murine PD-1 3.944, while Kd of αPD-1 to human PD-1 was 2.599, murine PD-1 2.766. We transfected hPD-1 into Jurkat cells (Jurkat-PD-1 cell) in order to mimic PD-1^+^ T cells. Within the cell level, we found that IC50 of αPD-1-(iRGD)_2_ to Jurkat-PD-1 was about 0.04 μg/ml while that of αPD-1 was 0.06 μg/ml (**Figure 2d**). Flow cytometry revealed that αPD-1-(iRGD)_2_ attained stronger affinity to HGC27, NCI-N87, MFC and B16F10 cancer cell lines than unmodified αPD-1 (**Figure 1g-j**). Competitive inhibitors (free iRGD and αNRP1) at least partially abrogated the affinity of αPD-1-(iRGD)_2_ towards cancer cell lines. However, both αPD-1 and αPD-1-(iRGD)_2_ exhibited little binding to HBE, 293T and GES-1 normal cell lines implying attachment of iRGD would not increase the accumulation of αPD-1 in normal tissue (**Figure 1k-m**).

**Figure 1.**
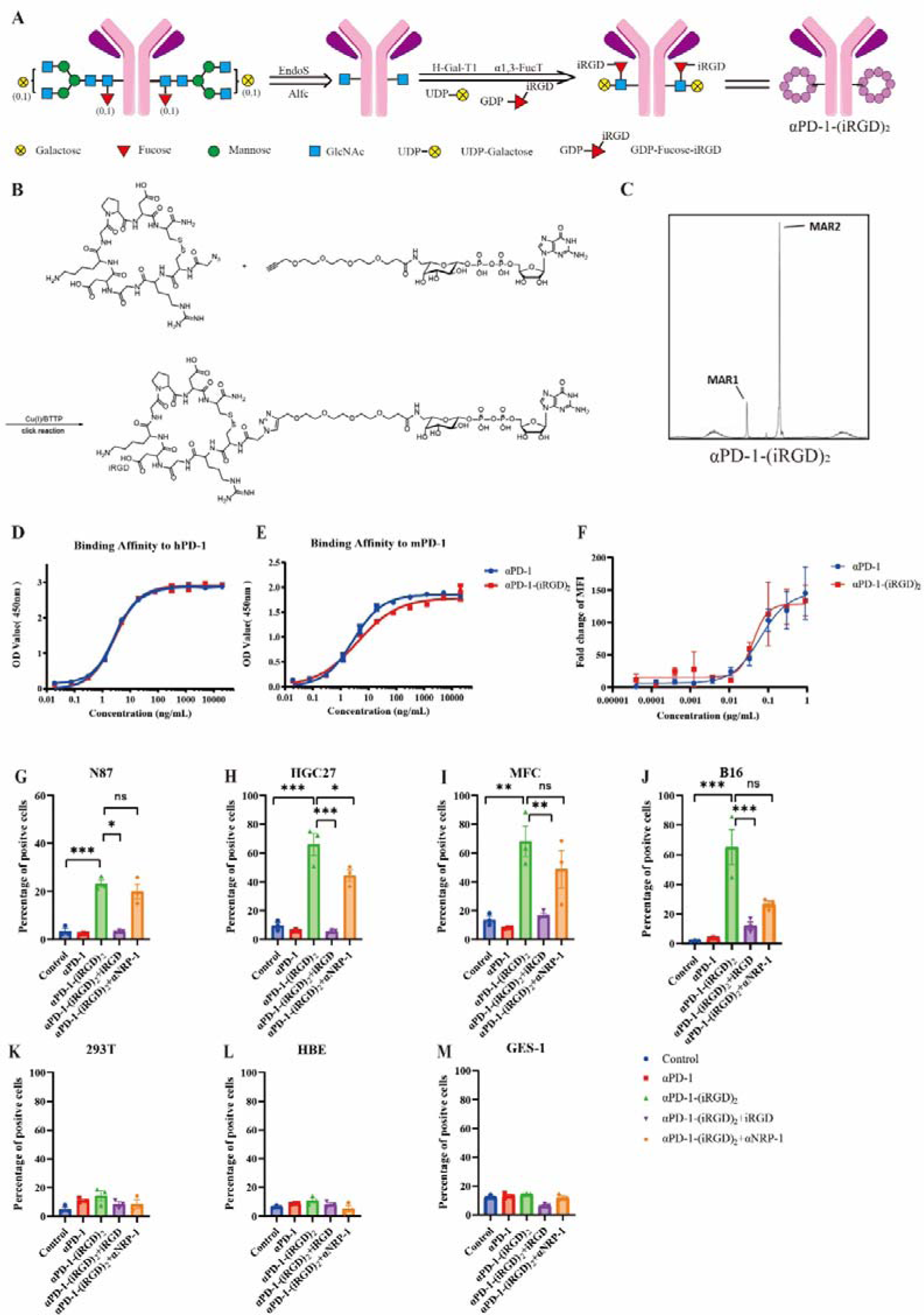
Synthesis and characterization of αPD-1-(iRGD)_2_ (A) The pattern diagram of the structure and chemical synthesis procedures of αPD-1-(iRGD)_2_. **(B)** Production of cor substrate GDP-Fucose-iRGD. **(C)** ESI-MS characterization of αPD-1-(iRGD)_2_. **(D-E)** The binding affinity of αPD-1-(iRGD)_2_ and unmodified antibody towards human (D) and murine (E) PD-1 protein by ELISA. **(F)** MFI fold change of Jurkat-PD-1 cells after being incubated with αPD-1 or αPD-1-(iRGD)_2_ for 1h. Data represent mean ± s.e.m.; n = 3. **(G-M)** Flow cytometric analysis of changes in relative averaged fluorescence intensities of cancer cell lines (N87, HGC27, MFC, B16) and normal cell lines (HBE, 293T, GES-1) cocultured with 10 μg/ml αPD-1-(iRGD)_2_ or 10 μg/ml αPD-1. The concentration of free iRGD was 100 μg/ml, and αNRP1 was 15 μg/ml. Data represent mean ± s.e.m.; n = 3.

**Figure 2.**
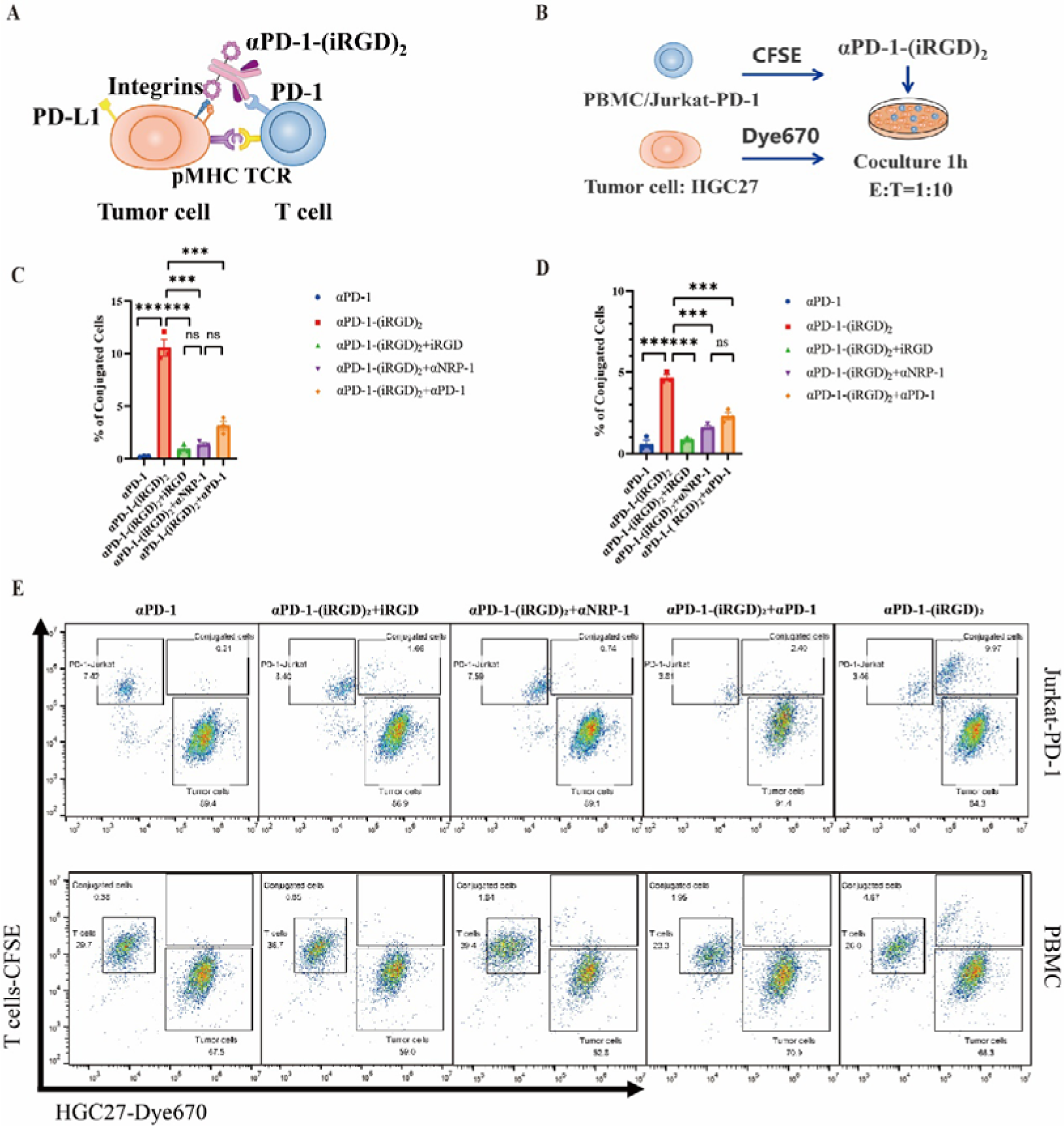
αPD-1-(iRGD)2 engages tumor cells and T cells (A) Diagram of αPD-1-(iRGD)_2_ engaging tumor cells and T cells. **(B)** Scheme of conjugate formation assay. **(C)** Percentage of CFSE and Dye670 double positive cells in HGC27 cells incubated with Jurkat-PD-1 cells at an E: T of 1:10. **(D)** The percentages of CFSE and Dye670 double positive cells in HGC27 cells incubated with PBMCs at an E: T of 1:10. **(E)** Representative flow cytometry results of (C-D). Data represent mean ± s.e.m.; n = 3. Student’s t test. n.s, not significant; *P < 0.5; **P < 0.01; ***P < 0.001.

### αPD-1-(iRGD)_2_ engages PD-1^+^ T cells and tumor cells

Due to the dual targeting capacity of αPD-1-(iRGD)_2_ to PD-1 and iRGD receptors, we first examined whether αPD-1-(iRGD)_2_ might engage T cells and tumor cells (**Figure 2a**). Considering that under physiological conditions limited T cells express PD-1, we apply CD3/CD28 beads and IL2, IL7, IL15 to induce exhaustion of PBMC and OT-I cells. PD-1 Expression of PBMC, OT-I and Jurkat-PD-1 were checked before subsequent assays (Figure S2-4). Afterwards, we conducted conjugate formation assay at an effector target ratio (E: T) of 1:10 with both Jurkat-PD-1 cells, exhausted PBMC from *HLA-A*2402*^+^ healthy donor, and *HLA-A*2402*^+^ HGC27 cells. Flow cytometry exhibited significantly increased cell engagement between PD-1^+^ T cells (OT-I, Jurkat-PD-1, PBMC) and tumor cells in αPD-1-(iRGD)_2_ group, compared with αPD-1 along (**Figure 2c-d**, Figure S5). What’s more, the engagement of T cells and tumor cells was competitively inhibited by iRGD, αNRP-1 and αPD-1. Fluorescence imaging of 5, 6-carboxyfluorescein diacetate, succinimidyl ester (CFSE) labeled HGC27 cells and CellTracker™ Deep Red Dye-labeled T cells cocultured with different agents visually displayed the engagement of cells mediated by αPD-1-(iRGD)_2_ (Figure S6). After that, we focused on whether αPD-1-(iRGD)_2_ could mediate better tumor killing efficacy. In both monolayer culture system and multicellular spheroids (MCS) system, αPD-1-(iRGD)_2_ effectively promoted the cytotoxicity of MHC-matched T cells against cancer cells in a concentration-dependent manner (**Figure 3a-b**). Such killing enhancement effect could also be inhibited by excessive dissociative iRGD, αNRP-1 and αPD-1 (**Figure 3c**). We further tested the capacity of αPD-1-(iRGD)_2_ to activate T cells. After coculturing, αPD-1-(iRGD)_2_ significantly elevated the surface expression of activation markers (CD69 and CD25) and cytotoxicity markers (GZMB and IFNγ) on T cells (**Figure 3d-g**). Similarly, αPD-1-(iRGD)_2_ magnified the antitumor efficacy of OT-I cells to B16-OVA cells. In the presence of B16-OVA cells, αPD-1-(iRGD)_2_ upgraded the level of CD25, CD69, GZMB and IFNγ while competitive inhibitors could partially abrogate such effect (Figure S7). This is an interesting finding, as it supports the idea that apart from penetration enhancement, αPD-1-(iRGD)_2_ can be considered a novel format of a bispecific T cell engager (BiTE) that does not target CD3.

**Figure 3.**
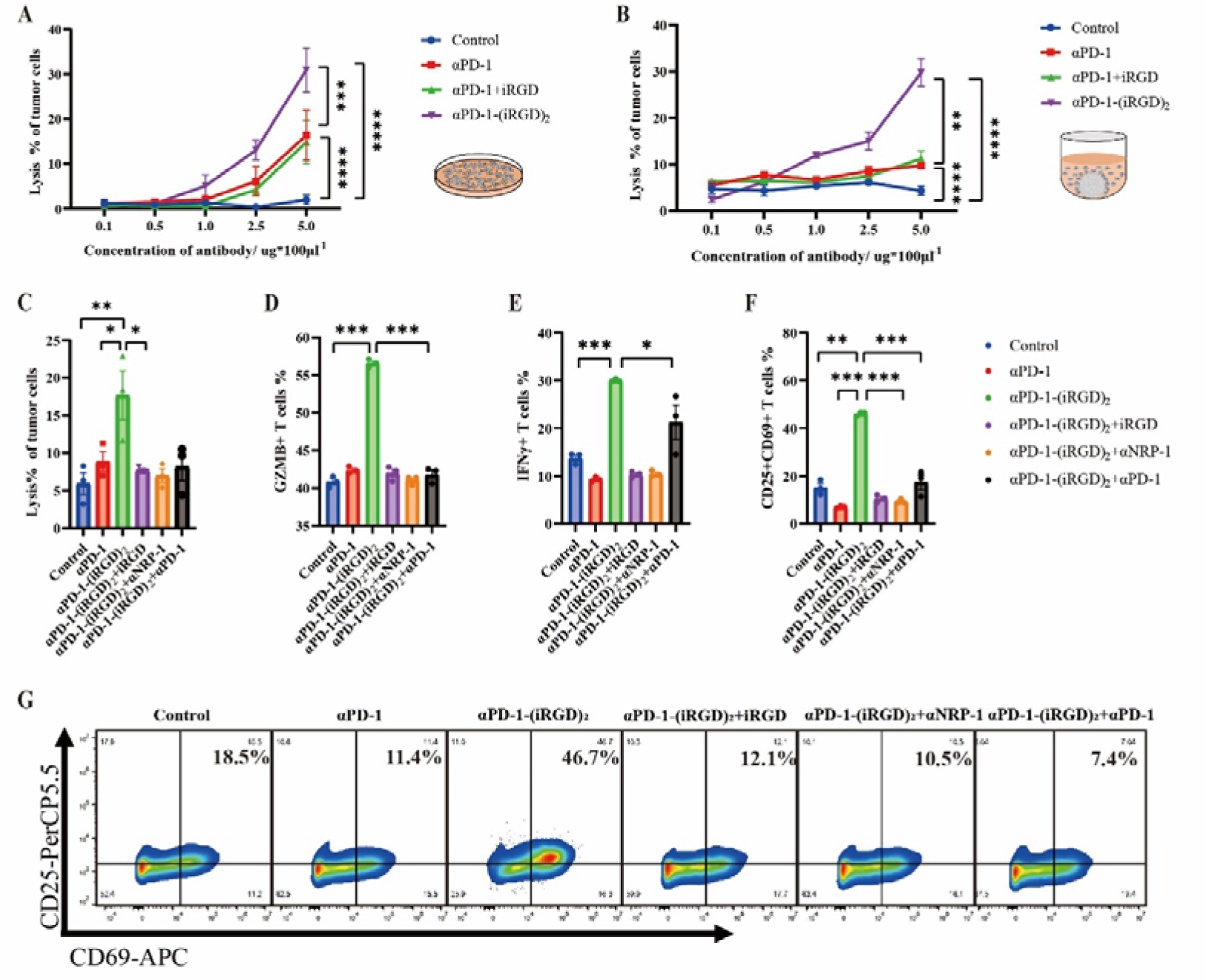
αPD-1-(iRGD)2 promotes T cell activation and tumor elimination (A) Lysis % of monolayer HGC27 cells coculturing with *HLA-A*2402*^+^ PBMC at an E: T rate of 10:1 under indicated antibody concentration. The concentration of free iRGD if added was 10μg/ml. **(B)** Lysis % of multicellular spheroids (MCSs) constructed with HGC27 cells coculturing with *HLA-A*2402*^+^ PBMC from healthy donor at an E: T rate of 10:1 under indicated antibody concentration. Concentration of free iRGD if added was 10μg/ml. **(C)** Lysis % of monolayer HGC27 cells coculturing with *HLA-A*2402*^+^ PBMC at an E: T rate of 10:1. The concentration of αPD-1, αPD-1-(iRGD)_2_ was 10μg/ml, iRGD was 100μg/ml, αNRP1 was 15μg/ml. Specially in αPD-1-(iRGD)_2_+αPD-1 group, the concentration of αPD-1 was 50μg/ml. **(D)** The expression of GZMB in PBMC cells under coculture conditions in (C). **(E)** The expression of IFNγ in PBMC cells under coculture conditions in (C). **(F)** The expression of CD25 and CD69 in PBMC cells under coculture conditions in (C). **(G)** Representative flow cytometry results of **(F).** Data represent mean ± s.e.m.; n = 3. Student’s t test. n.s, not significant; *P < 0.5; **P < 0.01; ***P < 0.001.

### αPD-1-(iRGD)_2_ promotes infiltration of PD-1^+^ T cells into tumor spheroids and syngeneic mouse tumors

In order to analyze the penetrability of αPD-1-(iRGD)_2_, we constructed MCSs with human gastric cancer cell line, HGC27 (**Figure 4a**). αPD-1-(iRGD)_2_-Cy5 infiltrated rapidly and robustly into the core of MCSs overtime, compared with αPD-1 alone or with different dose of free iRGD (**Figure 4c, e**). The high-does (100 μg/ml) free iRGD group failed to exhibit equivalent penetration to the αPD-1-(iRGD)2-Cy5 group even though it administered up to 10 times the amount of iRGD actually conjugated in αPD-1-(iRGD)_2_-Cy5. Apart from αPD-1-(iRGD)_2_-Cy5 itself, we further examined the capacity of αPD-1-(iRGD)_2_ to promote the infiltration of PD-1^+^ T cells (**Figure 4b**). αPD-1-(iRGD)_2_ extensively promoted CFSE labeled PD-1^+^ T cells entering the core of MCSs, while fluorescence signal of other groups was constrained to the periphery of the MCSs (**Figure 4d, f**). These results indicate that αPD-1-(iRGD)_2_ effectively penetrates MCSs and delivers T cells into the core area of MCSs.

**Figure 4.**
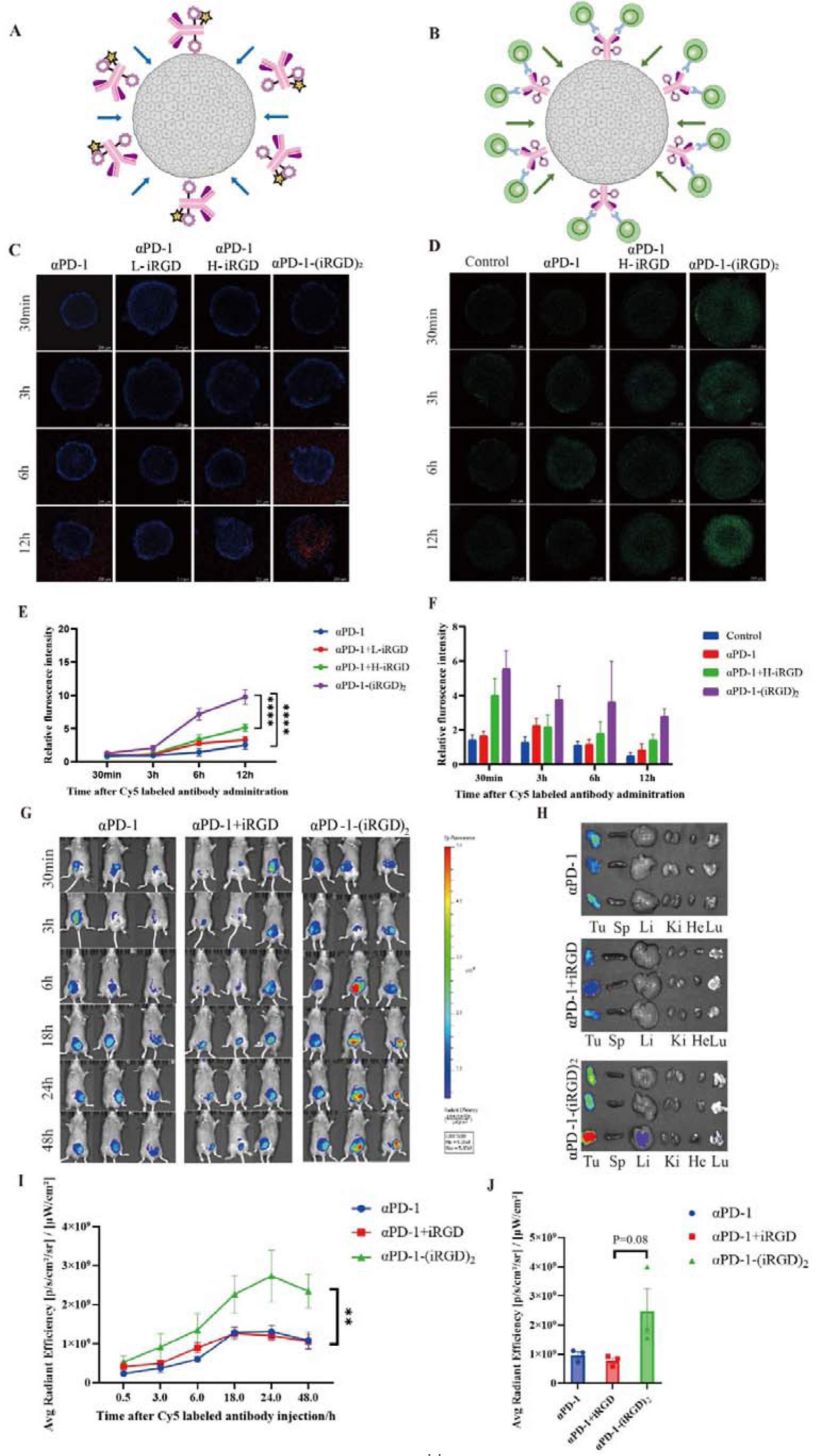
Penetrability and targetability of αPD-1-(iRGD)_2_ (A) Diagram of penetrability assay of Cy5 fluorescence labeled αPD-1-(iRGD)_2_. **(B)** Diagram of αPD-1-(iRGD)_2_ promoting CFSE labeled Jurkat-PD-1 cells infiltrating into HGC27 MCSs. **(C)**&**(D)** Confocal microscopy images showing penetration of Cy5-labeled antibody **(C)** and CFSE-labeled Jurkat-PD-1 cells **(D)** in MCSs. Magnification, ×50; scale bar, 200 µm. **(E)** The relative fluorescence intensity of MCSs incubated with Cy5 labeled antibody or control agents under indicated periods. **(F)** The relative fluorescence intensity of MCSs cocultured with CFSE-labeled Jurkat-PD-1 cells and αPD-1-(iRGD)_2_ or control agents under indicated periods. **(G)** Bioluminescence images of Cy5-labeled antibody at intervals after intraperitoneal injection to mice inoculated with MFC cells 10 days ago. **(H)** Ex vivo images of tumor, liver, heart, kidney, lung and spleen at 24h after intraperitoneal injection. **(I)** Average radiant efficiency at tumor location of each mouse in **(G)**. **(J)** Average radiant efficiency of resected tumor bulk in **(G)** 48h after agents administered. Data represent mean ± s.e.m.; n = 3. One-way ANOVA test, **P < 0.01.

Afterwards, the penetrability of αPD-1-(iRGD)_2_ was examined in the murine homologous tumor model. Near-infrared imaging of the living body displayed rapid and intensive distribution of αPD-1-(iRGD)_2_-Cy5 at tumor bulk (**Figure 4g, i**). 48 hours after injection, tumor and main organ were resected to evaluate the fluorescence intensity. The average radiant efficacy of tumor bulk doubled compared with αPD-1(**Figure 4j**). No significant increase in organ absorption was observed (**Figure 4h**). These observations were consistent with the *in vitro* findings that αPD-1-(iRGD)_2_ gained potent affinity to tumor cells. Conclusively, iRGD conjugation improves MCS penetration and tumor nodule targetability of αPD-1.

### αPD-1-(iRGD)_2_ effectively reduces tumor growth in murine tumor models

Immune checkpoint inhibitors aim to mobilize the host’s immune system, rescuing T cells from exhausted status and reviving immune response against cancer cells(24). Therefore, we selected immune competent murine tumor models to evaluate the antitumor efficacy of αPD-1-(iRGD)_2_. In MFC subcutaneous tumor model, mice treated with αPD-1-(iRGD)_2_ exhibited delayed tumor growth and remarkably lighter tumor weight at the end point (**Figure 5a-d**). αPD-1 alone also inhibited the tumor growth to a certain degree, while αPD-1 combined with free iRGD showed little difference to the αPD-1 group. Furthermore, flow cytometry results revealed that αPD-1-(iRGD)_2_ recruited more CD8^+^ T cells into the TME than other groups (**Figure 5i**). No difference was observed in PD-1 expression on CD8^+^ TILs, which may be attributed to the steric effect of PD-1 antibody(25). However, as for biomarkers of T cell, αPD-1-(iRGD)_2_ significantly elevated the expression of IFNγ and GZMB in both CD8^+^ and CD4^+^ TILs, while αPD-1 alone or with free iRGD exhibited slight influence (**Figure 5j, k**). Similar trend was also observed in tumor draining lymph node (Figure S8). During the treatment, weight loss or other side effects were not observed (Figure S9a). At the end of the experiment, no abnormal damage was recorded in the main organs as indicated by H&E staining (Figure S9c). These results demonstrates that αPD-1-(iRGD)_2_ exhibits significant tumor suppression effects with few side effects in the mouse model.

**Figure 5.**
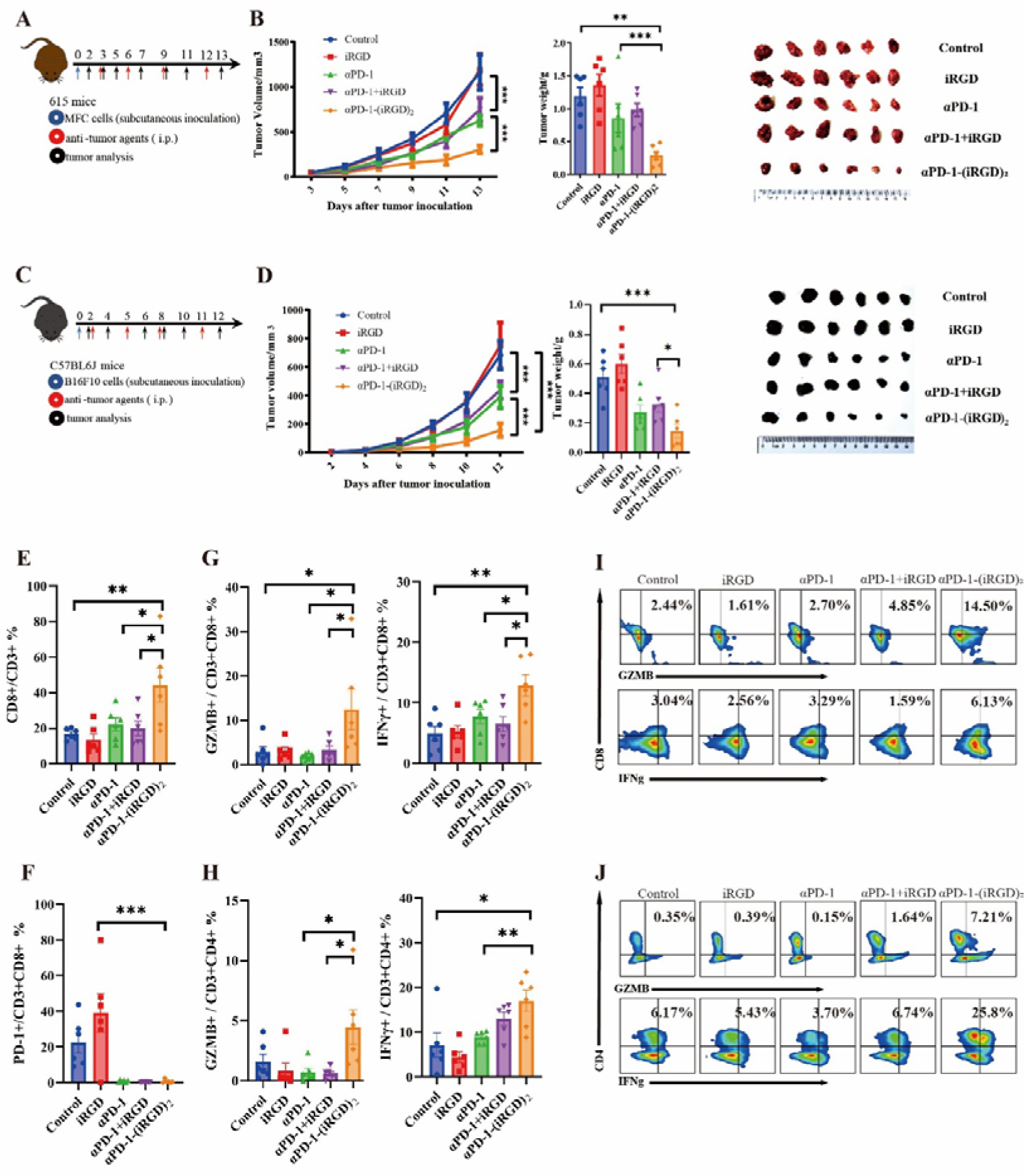
Antitumor efficacy of αPD-1-(iRGD)_2_ (A) Schematic of the treatment regimen in MFC mouse gastric tumor model. Briefly, mice were treated with 1 × 10^6^ MFC cells and injected intraperitoneally with PBS(100ul control), αPD-1(5mg/kg) alone or with free iRGD(1.25μg), αPD-1-(iRGD)_2_ (5 mg/kg) every three days. **(B)** Tumor growth profile and weight of tumors collected at the end point of **(A)**. **(C)** Schematic of the treatment regimen in B16F10 mouse melanoma model. Similarly, mice were subcutaneously inoculated with 1 × 10^5^B16F10 cells and injected intraperitoneally with PBS(100ul control), αPD-1(5mg/kg) alone or with free iRGD(1.25ug), αPD-1-(iRGD)_2_ (5 mg/kg) every three days. **(D)** Tumor growth profile and weight of tumors collected at the end point of **(C)**. **(E)** The abundance of CD8^+^ T cell from resected tumor bulk at the end point of **(A)**. **(F)** Flow cytometry quantification of PD-1 expression on CD8^+^ T cell from resected tumor bulk at the end point of **(A)**. **(G)** Flow cytometry quantification of GZMB and IFNγ expression on CD8^+^ T cells from resected tumor bulk at the end point of **(A)**. **(H)** Flow cytometry quantification of GZMB and IFNγ expression on CD4^+^ T cells from resected tumor bulk at the end point of **(A)**. **(I)** Representative flow cytometry results of GZMB and IFNγ expression on CD8^+^ T cells from resected tumor bulk at the end point of **(A)**. **(J)** Representative flow cytometry results of GZMB and IFN γ expression on CD4^+^ T cells from resected tumor bulk at the end point of **(A)**. Data represent mean ± s.e.m.; n = 6. One-way ANOVA; n.s, not significant; *P < 0.5; **P < 0.01; ***P < 0.001; ****P < 0.0001.

The antitumor efficacy of αPD-1-(iRGD)_2_ was also tested in B16F10 murine melanoma model, which was considered a “cold tumor” due to poor immune infiltration. Impressively, αPD-1-(iRGD)_2_ also postponed tumor growth and remarkably lightened tumor weight at the end point indicating that αPD-1-(iRGD)_2_ may have broad-spectrum antitumor effects (**Figure 5e-h**).

### αPD-1-(iRGD)_2_ remodels tumor immune microenvironment

We performed single cell RNA sequencing of CD45^+^ tumor infiltrating immune cells to further depict the immune landscape after the administration of αPD-1-(iRGD)_2_. We analyzed these CD45^+^ cells from different treatment groups using Uniform Manifold Approximation and Projection (UMAP) (**Figure 6a-c**). UMAP displayed 27 clusters including 5 cell types. According to specific cell markers, we defined clusters 4,7,10,11,12,15,18,23 into T cells (Trac, Trbc, Cd3d), clusters 6,9,19,22 as DC (Cd74, H2-Aa), clusters 0,2,6,8,14,16,21,24 as Macrophages (Adgre1, Itgax, Cd83), clusters 3,5,20 as Monocytes (Cd14), clusters 12,26 as NK cells (Nkg7, Klrb1c, Klrg1) (Figure S10). Consistent with our previous findings, the expression levels of several effector genes (*Gzma*, *Ifng*, *Ifngr1*, etc.), inflammatory cytokine receptors (*Il2ra*, *Il7r*, *Il12rb1*, etc.) and co-stimulatory factors (*Cd28*, *Cd27*, etc.) were upregulated in CD8^+^ TILs from mice that received αPD-1-(iRGD)_2_ (**Figure 6e**). Similar transcriptional profiles were also observed in CD4^+^ TILs (**Figure 6f**). One of the most notable changes in CD8^+^ T cells after αPD-1-(iRGD)_2_ administration was the upregulation of several stem and memory associated genes (*Lef1*, *Tcf7*, *Bach2*, *Ikzf2*)(26-28). The RNA levels of genes involved in migration and adhesion (*Ccr2*, *Cxcr3*, *S1pr1*, *Itgb1* and *Ly6c2*) also increased, indicating that αPD-1-(iRGD)_2_ recruited CD8^+^ T cells to infiltrate into TME.

**Figure 6.**
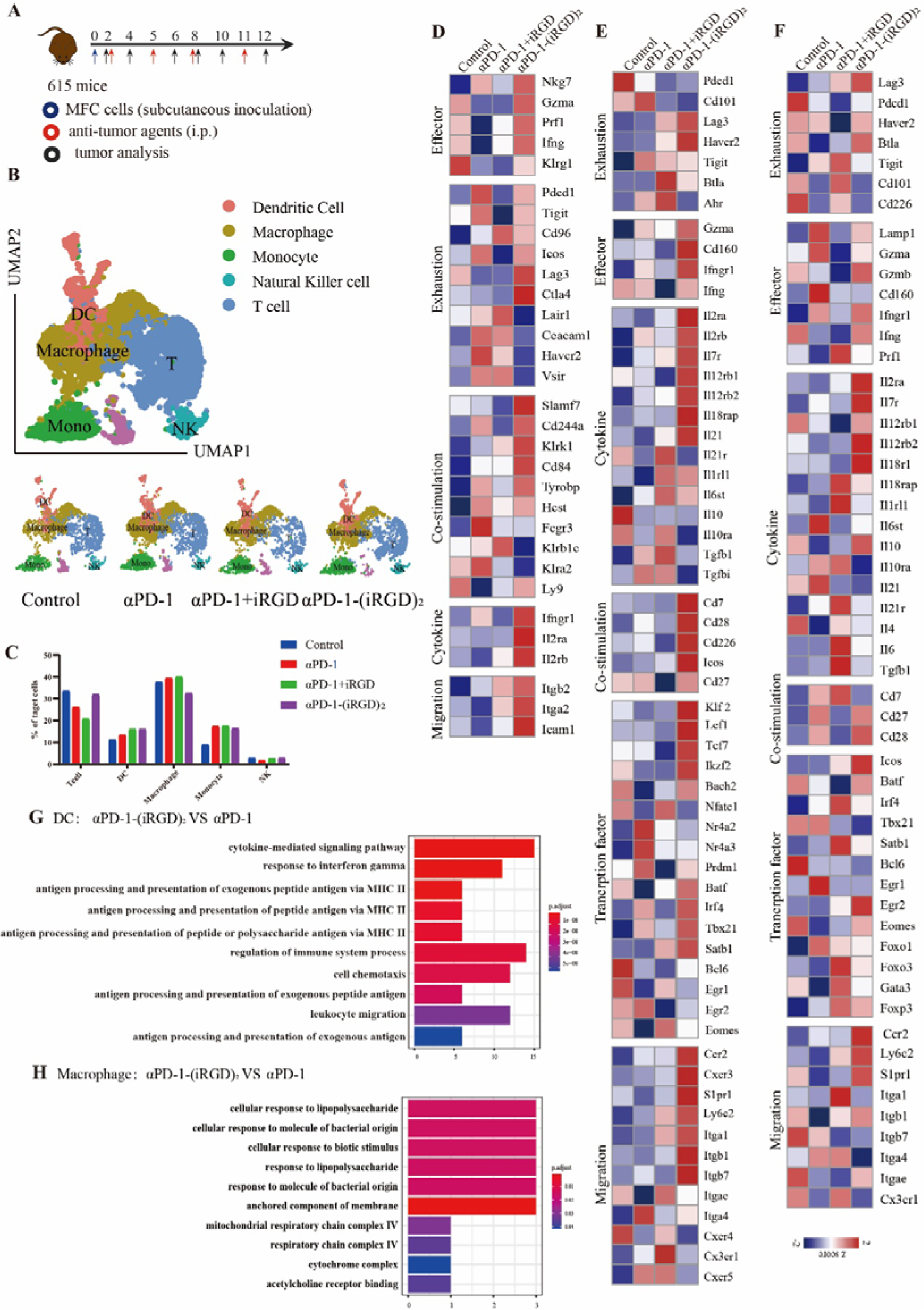
αPD-1-(iRGD)2 remodels tumor microenvironment (A) Schematics of the treatment regimen in MFC homograft models are shown. Briefly, mice bearing tumor burdens were treated with 1 × 10^6^ MFC cell and injected intraperitoneally with PBS(100μl control), αPD-1 (5mg/kg) alone or with free iRGD(50μg), αPD-1-(iRGD)_2_ (5 mg/kg) every three days. **(B)** Two-dimensional (2D) UMAP visualization of CD45^+^ tumor infiltrating immune cells coloured according to subset (upper) and specific treatment group (down). **(C)** The ratio of each subset within the CD45^+^ tumor infiltrating immune cells is depicted in **(B)**. **(D)** Heatmap of average relative expression of selected genes in NK cells across treatment groups within CD45^+^ tumor infiltrating immune cells. **(E)**&**(F)** Heatmap of average relative expression of selected genes in CD8^+^ **(E)** and CD4^+^ **(F)** T cells across treatment groups within CD45^+^ tumor infiltrating immune cells depicted in **(B)**. **(G)** KEGG analysis of differentially expressed genes in DC betweenαPD-1 and αPD-1-(iRGD)_2_. **(H)** KEGG analysis of differentially expressed genes in macrophage between αPD-1 and αPD-1-(iRGD)_2_.

Apart from T cells, NK cells, DC and macrophages also participated in tumor suppression of αPD-1 directly or indirectly(29-31). In mice treated with αPD-1-(iRGD)_2_, NK cells in TME expressed higher level of effector genes (*Nkg7*, *Gzma*, *Prf1*, *Klrg1*), co-stimulatory factors (*Slamf7*, *Cd244a*, *Klrk1*), migration genes (*Itgb2*, *Itga2*, *Icam1*), immune stimulatory cytokines and receptors (*Ifng*, *Ifngr1*, *Il2ra*, *Il2rb*) (**Figure 6d**). Kyoto Encyclopedia of Genes and Genomes (KEGG) analysis was performed using differentially expressed gene signatures of DCs and macrophages. The results showed an enrichment of response to IFNγ, antigen processing and presentation in DC from mice treated with αPD-1-(iRGD)_2_ (**Figure 6g**). Differentially expressed genes of macrophage enriched in cellular response to lipopolysaccharide and biotic stimulus as well (**Figure 6h**). These observations indicated αPD-1-(iRGD)_2_ simultaneously ameliorated the function of NK, DC and macrophage. In conclusion, αPD-1-(iRGD)_2_ profoundly remodulates the immune suppressive TME thus facilitating the cytotoxicity of CD8^+^ T cells.

To gain further insights into how the differentiation program of CD8^+^ T cells was altered by αPD-1-(iRGD)_2_ versus PD-1 monotherapy, we reperformed UMAP analysis of predefined T cells (**Figure 7a**). UMAP displayed 10 clusters and were annotated into 6 subtypes according to cell markers, including better effector CD8^+^ T cells (cluster 4), effector CD8^+^ T cells (cluster 3), intermediate exhausted CD8^+^ T cells (clusters 0,2,5), terminal exhausted CD8^+^ T cells (clusters 1,9), memory CD8^+^ T cells (cluster 6), CD4^+^ T cells (clusters 7, 8) (**Figure 7b-c**). Impressively, the abundance of a unique subset of CD8^+^ T cells (better effector CD8^+^ T cells), which featured in the expression of stem and memory associated genes (*Tcf7*, *Il7r*, *Lef1*, *Bach2*) and intermediate expression of effector genes (*Gzma*, *Gzmb*, *Ifng*), expanded in TILs of αPD-1-(iRGD)_2_ treated mice (**Figure 7d**). In addition, this population of CD8^+^ TILs also expressed low levels of transcription for *PD-1*, *Lag3*, *Havcr2* (*TIM-3*) and *Entpd1* in line with a non-exhausted profile. According to previous work that PD-1-cis IL-2R agonism yields better effectors from stem-like CD8+ T cells, we identified this subset of CD8^+^ TILs as “better effector” T cells underlining their highly functional effector profile and lower degree of exhaustion(32, 33). The expansion of better effector CD8^+^ T cells implies that αPD-1-(iRGD)_2_ reinforceed the effector function of TILs while avoiding exhaustion and maintained stem and memory characteristics of CD8^+^ TILs.

**Figure 7.**
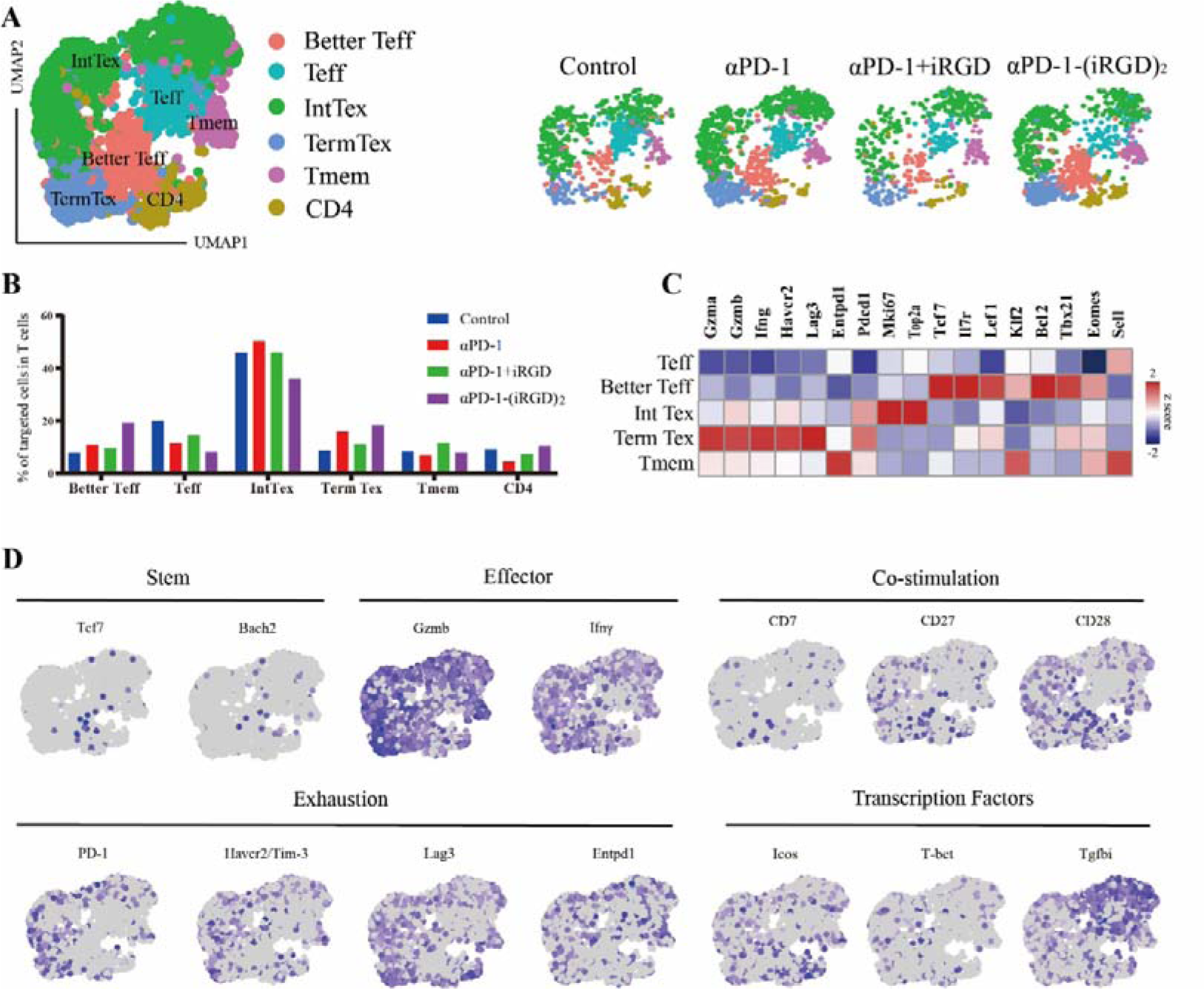
αPD-1-(iRGD)2 expands a unique population of better effector CD8+ TILs (A) Two-dimensional (2D) UMAP visualization of CD3^+^ TILs coloured according to subset (Left) and specific treatment group (Right). **(B)** Ratio of each subset within the CD3^+^ tumor infiltrating T cells depicted in **(A)**. **(C)** Heatmap of normalized expression of several representative genes within the clusters of CD3^+^ TILs depicted in **(A)**. **(D)** Normalized expression of several representative genes within the six clusters of CD3^+^ TILs.

**Figure 8.**
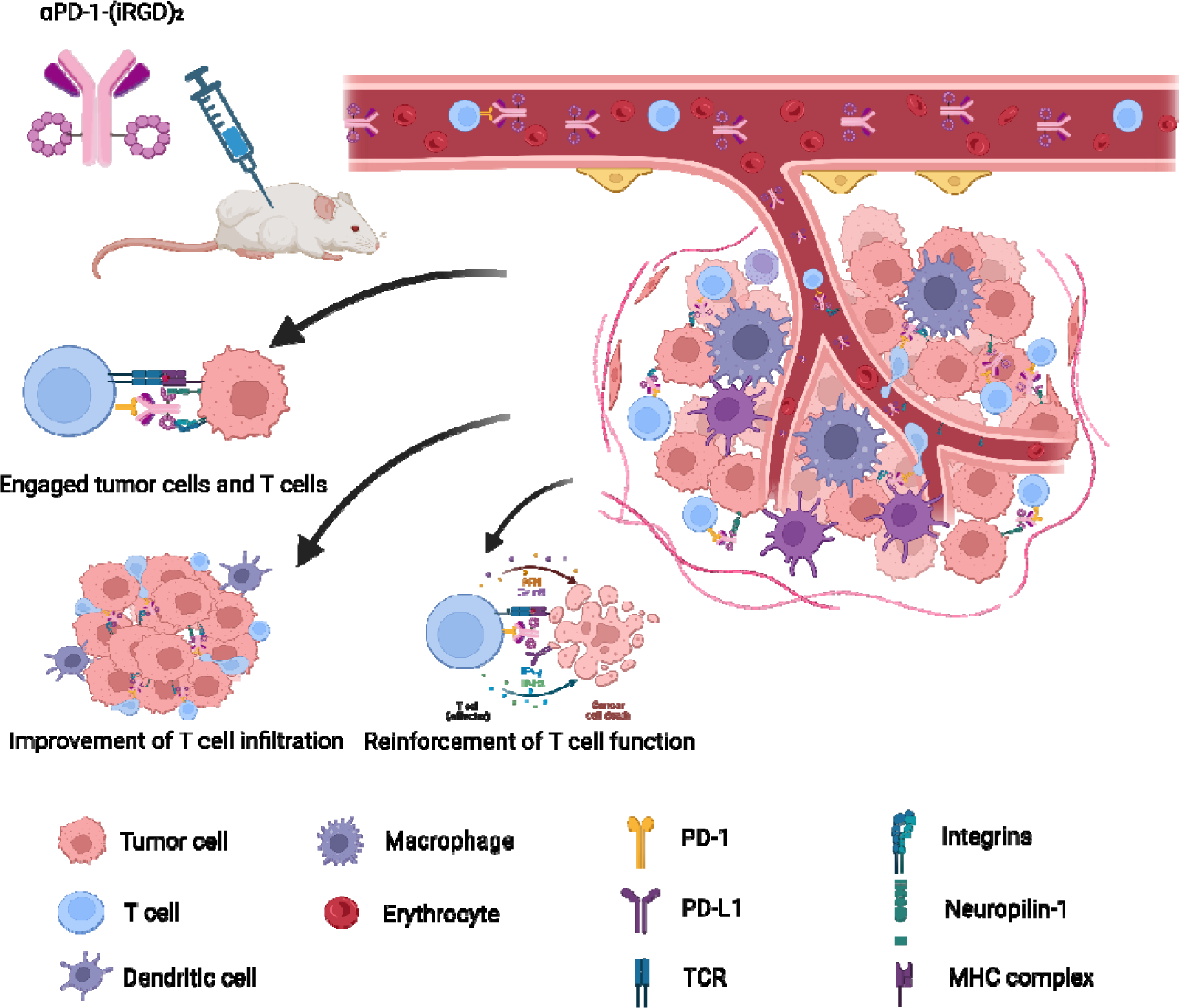
Schematic diagram of antitumor effect of αPD-1-(iRGD)_2_. A proposed model for the multidimentional antitumor efficacy of αPD-1-(iRGD)_2_, including engagement of T cells and tumor cells, improvement of T cell infiltration, and replenishiment of T cell function.

### Construction and evaluation of murine αPD-1-(iRGD)_2_

Although CS1003 binds both human and murine PD-1, its humanized Fc domain may cause unpredictable influence on the antitumor efficacy of αPD-1-(iRGD)_2_. In an effort to exclude the incompatible Fc domain and examine the suitability of LacNAc-conju to murine antibodies, we constructed αPD-1-(iRGD)_2_ from murine αPD-1 (G4C2, TopAlliance). ESI-MS analysis confirmed that the product possessed an expected molecular weight, while ELISA results exhibited that αPD-1-(iRGD)_2_ sustained its affinity to murine PD-1 (Figure S11a-b). After that, the tumor suppression function of αPD-1-(iRGD)_2_ was evaluated in the murine gastric cancer tumor model. Murine αPD-1-(iRGD)_2_ displayed superior antitumor efficacy than αPD-1 monotherapy (Figure S11c-d).

Collectively, our results further prove that iRGD conjugate mediated valid improvement of antitumor effect of αPD-1.

## Discussion

ICI, represented by PD-1 antibodies, has emerged as standard regimen for several solid tumors(34). However, the ORR of ICI remains limited which restrains its broader application. Inefficient biodistribution of therapeutic agents and insufficient infiltration of T cells within tumor microenvironment (TME) partly account for the restrained clinical response. Therefore, methods to enhance the therapeutic effects of ICI are intensively required. iRGD peptide has been proved to boost the penetration of therapeutic agents into tumor bulk via successively binding to integrins and NRP-1(17). In an open-label, multicenter, phase I clinical study, iRGD peptide in combination with chemotherapeutics preliminarily exhibited superior antitumor efficacy in metastatic pancreatic ductal adenocarcinoma(35). Our team previously anchored iRGD to the surface of T cells thus facilitating the infiltration of T cells into MCSs and tumor bulk(18). We also designed anti-CD3-iRGD, a bifunctional agent that possesses dual affinity to CD3 and iRGD receptors and resembles BiTEs which could provide an extra activation signal to further strengthen the antitumor effect(19). However, anti-CD3 based BiTEs have several drawbacks, including recruitment of counterproductive CD3^+^ T cell subsets, the release of systemic cytokines, the expansion of immune checkpoint molecules, the presence of an immunosuppressive TME, tumor antigen loss or escape, on-target off-tumor toxicity and suboptimal potency(36). While almost all T cells express CD3, tumor-specific CD8^+^ TILs express high levels of PD-1 and are temporarily functionally impaired(37). Therefore, we chose αPD-1 as modification object. αPD-1-(iRGD)_2_ mainly acts on PD-1^+^ TILs and avoids non-selective T cell activation and excessive TCR signaling. Meanwhile, in this work, we found that αPD-1-(iRGD)_2_ not only facilitated the penetration of T cells but also engaged T cells and tumor cells due to its dual affinity. In monolayer coculture tumor killing assay, where penetration related factors were ruled out, αPD-1-(iRGD)_2_ still significantly enhanced T cell cytotoxicity. Besides, competitive inhibitors including free iRGD, αNRP1 and αPD-1 abrogated the engagement formation, tumor elimination and T cell activation mediated by αPD-1-(iRGD)_2_ to a certain degree. Especially, after exhaustion inducement with CD3/CD28 beads and interleukins, T cells abundantly expresses PD-1 and response mildly to αPD-1 alone. However, αPD-1-(iRGD)_2_ could still enhance tumor elimination and activation marker expression. Collectively, we suppose that αPD-1-(iRGD)_2_ may not only accord T cells with superior penetrability but also bridge TILs and Tumor cells to unleash more antitumor potential.

Unlike anti-CD3-iRGD generated by plasmid construction and expression, we performed site-specific modification of both human and murine αPD-1 via glycoengineering platform. Therefore, αPD-1-(iRGD)_2_ preserves the natural characteristics of antibodies. Due to its larger molecular size and FcRn mediated recycling processes, αPD-1-(iRGD)_2_ with an Fc region might have a longer half-life in circulation(38). Besides, αPD-1-(iRGD)_2_ is more convenient to purify and displays increased solubility and stability. More importantly, it may have greater clinical translational potential for retaining a diverse range of Fc-mediated effector functions(39).

Additionally, the dose of free iRGD (1.25ug) in the αPD-1+iRGD group was equivalent to the absolute scale of iRGD conjugated to αPD-1 in our experiment. Under such circumstances, the αPD-1+iRGD group did not exhibit improvement of antitumor efficacy over the αPD-1 group. Sugahara et al. disclosed that Nab-paclitaxel (ABX) conjugated with iRGD peptide had comparable antitumor efficacy to ABX combined with free iRGD, when iRGD was administered at the dose of 4μmol/kg every other day(17). However, previous researchers found the half-life of free iRGD was limited to minutes, which requires frequent administration. Meanwhile, high dose iRGD and αPD-1 combination could not mediate engagement of T cells and tumor cells. Summarily, conjugation generated unique BiTE-like antitumor mechanism, remarkably reduced the required dose of iRGD and allowed for longer injection interval thus improving treatment compliance and feasibility.

Last but not least, αPD-1-(iRGD)_2_ reinforces CD8^+^ T cells and expands a unique population of “better effectors”. PD-1^+^TCF-1^+^CD8^+^ T cells have emerged as important determinants of the immune response in chronic infections and cancer, with their abundance critical to the success of cancer immunotherapies(40, 41). In our work, after the administration of αPD-1-(iRGD)_2_, abundance of a subset of T cells, expressing stem and memory associated genes, intermediate effector genes and low levels of exhaustion markers, increased. According to previous studies, we defined them as “better effectors”(32). We speculate that the dual binding affinity of αPD-1-(iRGD)_2_ reinforces the effector function of TILs while avoiding exhaustion and maintained stem and memory characteristics of CD8^+^ TILs. The receptors of iRGD, including integrin αv and NRP1, have been reported to regulate antitumor CD8^+^ T cell immunity and response to PD-1 blockade(42). In addition, iRGD commands were reported to be molecular degraders for integrin-facilitated targeted protein degradation(43). Further exploration is required to clarify the modulation of integrin αV and NRP1 by iRGD.

However, there still remain several limitations in our work. The mechanism of αPD-1-(iRGD)_2_ mediating the expansion of “better effector” T cells with stem and memory phenotype needs to be further explored. Pharmacokinetics characteristics and further safety evaluation are required before clinical translation.

In the past decade, immunotherapy represented by PD-1/PD-L1 blockade has revolutionized the standard care for several types of tumors. Underperforming drug uptake and poor immune infiltration largely limited the clinical benefit. In this work, we performed site-specific modification to both human and murine αPD-1 via our one-step glycoengineering platform. αPD-1-(iRGD)_2_ can enhance therapeutic potential for cancer treatment by simultaneously engaging T cells and tumor cells, promoting T cell infiltration and expanding a unique population of “better effector” T cells. We believe αPD-1-(iRGD)_2_ provides a paradigm shift of anti-PD-1 based therapeutic approach with the ability to reshape the TME via a distinct combinatorial mechanism.

## Methods

### Synthesis of GDP-FAmP4PropP4-iRGD

To a solution of 500 uL GDP-FAm (100 mM) was added 1500 uL ddH_2_O, 500 uL NaHCO_3_ (200 mM), 1750 uL THF and 750 uL NHS-PEG_4_-Prop (Confluore) (50 mM in Tetrahydrofuran (THF)). The reaction was stirred at room temperature (r.t.) for 4 hours and monitored by thin layer chromatography (TLC). The solvent was removed under reduced pressure. The crude product was further purified through a Prep-HPLC system to give the desired product as a white solid (27.1 mg. 64%).

To a solution of 640 μL GDP-FAmP_4_Prop (50 mM) in ddH_2_O/DMSO (2048 μL/1920 μL), 64 μL CuSO_4_ (50 mM in ddH_2_O), 128 ul BTTP (50 mM in ddH_2_O), 1280 μL N_3_-iRGD (Nanjing YuanPeptide Biotech Ltd.) (50 mM in DMSO), and 320 μL ascorbate (50 mM in ddH2O) were added. The reaction as allowed for stirring at r.t. for 12 h and monitored by TLC. The crude product was further purified through a Prep-HPLC system to give the product as a white solid (38.4 mg, 64%). HRMS (ESI-) calcd for C_63_H_103_N_23_O_34_P_2_S_2_ (M-2H^+^)/2 936.7905, found 936.7932.

### Cloning, expression and purification of bovine β1,4-GalT1(Y289L), EndoS(Streptococcus pyogenes endoglycosidase S) and Alfc (Lactobacillus casei α-1,6-fucosidase)

The cloning, expression and purification of bovine β1,4-GalT1(Y289L), EndoS and AlfC were performed according to the reported procedure by Qasba, P. K et al., by Wang L., et al. and by Wu P. respectively(44-46).

### Cloning, expression and purification of *Helicobacter pylori* α1,3 fucosyltrasferase

Genes encoding the helicobacter pylori α1,3 fucosyltrasferase were synthesized and subcloned into a pET24b vector at NdeI and BamHI by Genscript. E. coli BL21(DE3) transformed with the plasmids were cultured at 37 °C in LB with 50 μg/mL kanamycin until OD600 = 0.6-0.8. IPTG was added to a final concentration of 0.2mM and protein expression was induced for sixteen hours at 25 °C. The cells were harvested by centrifugation and resuspended in lysis buffer (25 mM Tris pH 7.5, 500 mM NaCl, 20mM imidazole and 1 mM PMSF). Cells were lysed by sonication and the clarified supernatant was purified on Ni-NTA agarose (GE Health) following the manufacturer’s instructions. Fractions that were >90% purity, as judged by SDS-PAGE, were consolidated and dialyzed against Tris-buffered saline (25 mM Tris pH 7.5, 150 mM NaCl).

### Cloning, expression and purification of αPD-1

The sequence of αPD-1 antibody light chain and heavy chain were referenced to the patent (CN202210202373.1). The gene encoding the light chain and the heavy chain of αPD-1 were synthesized and clone into a PPT5 vector respectively by Genescript. Then, FreeStyle 293F cells were grown to a density of ∼2.5×10^6^ cells/ml and transfected by direct addition of 0.37 μg/ml and 0.66 μg/ml of the light chain and heavy chain expression plasmid DNA, and 2.2 μg/ml polyethylenimine (linear 25 kDa PEI, Polysciences, Inc, Warrington, PA) to the suspension cultures. The cultures were diluted 1:1 with Freestyle 293 expression medium containing 4.4 mM valproic acid (2.2 mM final) 24 h after transfection, and protein production was continued for another 4–5 d at 37 [. After protein production, the antibodies were purified through the protein A agarose following the manufacturer’s instructions.

“One-pot” synthesis of αPD-1-(Galβ1,4) GlcNAc-FAmP4PropP4-iRGD

αPD-1 (8 mg/mL) were incubated with EndoS (0.05 mg/mL) and Alfc (1.5mg/mL) in 50 mM Tris-HCl buffer (pH 7.5) at 30°C. After 24 h, the reaction mixtures were added with UDP-Gal (final concentration 5 mM), bovine β1,4-GalT1(Y289L) (final concentration 0.3 mg/mL), GDP-FAmP4PropP4-iRGD (final concentration 2 mM) and Hp1,3-FucT (final concentration 0.5 mg/mL), MgCl_2_ (final concentration 10 mM) and MnCl_2_ (final concentration 5 mM) followed by incubating at 30°C for 48 h. The modified antibody was purified with protein A resin to give the αPD-1-(Galβ1,4) GlcNAc-FAmP4PropP4-iRGD (found as 148833.00 Da, MAR2) conjugates.

### Binding affinity assay

Recombinant PD-1 extracellular domains (PD-1, novoprotein) was diluted to a final concentration of 250 ng/mL with coating buffer and plated on 96-well plates (100 μL/well) at 4 °C for overnight. After removing the coating solution, the plates were blocked with 3% (v/v) bovine serum albumin in PBS for 2 h at 37[. After washing with PBST (PBS containing 0.03% tween-20) for 3 times, αPD-1 and αPD-1-(Galβ1,4)GlcNAc-FAmP4PropP4-iRGD were added to PBST (with 1% (v/v) bovine serum albumin in PBS) to a series of final concentrations (3000 ng/mL, 1000 ng/mL, 333.33 ng/mL, 111.11 ng/mL, 37.04 ng/mL, 12.35 ng/mL, 4.12 ng/mL, 1.37 ng/mL, 0.46 ng/mL, 0.15 ng/mL, 0.05 ng/mL, 0 ng/mL) and added to the plates respectively. After incubating for 1.5 h, the plates were washed 3 times with PBST, then horseradish peroxidase (HRP)-conjugated goat anti-human IgG antibody was added to each well and incubated for 1 h at 37[. Finally, each well was washed with PBST for 3 times, and then tetramethyl benzidine substrate was cotreated to produce color for visualization. The reaction in each well was stopped by adding 100 μL of 3 M HCl after 15 min of incubation. The absorbance was read at 450 nm on a Synergy^TM^ LX plate reader.

### Intact protein mass analysis

For LC-MS analysis, the purified proteins were analyzed on an Xevo G2-XS QTOF MS System (Waters Corporation) equipped with an electrospray ionization (ESI) source in conjunction with Waters Acuqity UPLC I-Class plus. Separation and desalting were carried out on a waters ACQUITY UPLC Protein BEH C4 Column (300 Å, 1.7 µm, 2.1 mm x 100 mm). Mobile phase A was 0.1% formic acid in water and mobile phase B was acetonitrile with 0.1% formic acid. A constant flow rate of 0.2 ml/min was used. Data were analyzed using Waters Unify software. Mass spectral deconvolution was performed using a Unify software (version 1.9.4, Waters Corporation).

### Cell lines

Human gastric cancer cell lines HGC27 and NCI-N87, human normal tissue cell lines HBE, GES-1 and 293T, a human leukemic T-cell line cell Jurkat, a mouse gastric cancer cell line MFC and melanoma cell lines B16F10, B16F10-OVA were purchased from the Cell Bank of Shanghai Institute of Biochemistry and Cell Biology (Shanghai, China) in 2018. Jurkat cells were transduced with lentiviruses generated from PLV-EF1a-PD-1. All of the cells mentioned above were cultured in Roswell Park Memorial Institute (RPMI) 1640 (Corning) supplemented with 10% fetal bovin serum (FBS, Gibco), 1% penicillin/streptomycin (Beyotime) at 37°C and 5% CO_2_. Cells were regularly tested for Mycoplasma with PCR method. The most recent cell line authentication was in September 2020 by short tandem repeat (STR) analysis.

### OT-I cells acquisition and culture

Spleens of OT-I mice were resected and mechanically dissociated. Single cell suspensions were obtained through 40-µm nylon cell strainers (Biosharp). Splenocytes from OT-I mice were cultured at about 1 × 10^6^ cells/ml in 24 well-plates with RPMI (Thermo Fisher) along with 10% FBS (Thermo Fisher), 1% penicillin/streptomycin (Thermo Fisher), 2mM L-glutamin (Thermo Fisher), 50μM β-mecaptoethanol (Thermo Fisher), 10mM HEPES (Thermo Fisher) and IL-2 (100U/ml, Thermo Fisher) in the presence of OVA_257-264_ peptide (0.1nM) at 37°C and 5% CO_2_. 3 days later, activated cells were washed 3 times with RPMI 1640 and resuspended in T25 flasks at 1 × 10^5^ cell/ml in the presence of IL-2(100U/ml), IL-7, IL-15(10ng/ml, Thermo Fisher).

### Mice

615-line mice were purchased from Institute of Hematology, Chinese Academy of Medical Sciences (Tianjin, China). C57BL/6 mice were purchased from Gem Pharmatech Co., Ltd. (Nanjing, China). All animals in our study were raised in the pathogen-free animal facilities at Nanjing University Medical School Affiliated Drum Tower Hospital (Nanjing, China). Six-to eight-week-old female or male mice were randomized based on age and weight and used for all *in vivo* experiments.

### Penetrability analysis of αPD-1-(iRGD)_2_ in MCSs

MCSs were generated with HGC27 cells as previously described(19). Subcircular MCSs with a diameter of around 500μm were selected for this study. αPD-1-(iRGD)_2_-Cy5 (10μg/ml) or αPD-1(10μg/ml) along or αPD-1(10μg/ml) with free iRGD(10μg/ml in L-iRGD, 100μg/ml in H-iRGD) were cocultured with MCSs for 24 hours at 37[. In cell penetrating assay, 1×10^6^ Jurkat-PD-1 cells were labeled with CFSE (Abcam, Cambridge, UK). After labeling, Jurkat-PD-1 cells along with αPD-1-(iRGD)_2_ (10μg/ml) or αPD-1(10μg/ml) along or αPD-1(10μg/ml) with free iRGD(100μg/ml), were cocultured with MCSs for 24 hours at 37[. Then, the MCSs were washed with PBS and fixed with 4% paraformaldehyde, and imaged using a ZEN 710 confocal microscope (Zeiss, Jena, Germany). Images were acquired close to mid-height of the spheroids. Fluorescence intensity was calculated with Leica Application Suite X (LAS X).

### *In vivo* real time near-infrared fluorescence imaging of Cy5 labeled αPD-1-(iRGD)_2_

Near infrared live body imaging was used to trace the distribution of αPD-1-(iRGD)_2_-Cy5 or αPD-1. 50ug αPD-1-(iRGD)_2_-Cy5, or αPD-1-Cy5 along or with 1μg free iRGD were injected i.p. into MFC tumor bearing mice with an average tumor volume of 200mm^3^. After anesthesia, mice were scanned with CRi Maestro *In Vivo* Imaging System (Cambridge Research & Instrumentation, Massachusetts, USA) at different time points after injection. 48h after agents’ administration, mice were humanely executed and tumor tissues and main organs were resected. The *In vitro* average radiant efficiency of tumor and main organ tissues were also scanned with CRi Maestro *In Vivo* Imaging System (Cambridge Research & Instrumentation, Massachusetts, USA).

### Cytotoxicity assay

*HLA-A*2402*^+^ PBMCs from healthy donor were collected via Ficoll density centrifugation. PBMCS were cultured in AIM-V medium (Gibco, USA) along with 10% FBS at 37°C and 5% CO_2_. 2 hours later, non-adherent T cells were collected and suspended in AIM-V medium (Gibco, USA) along with 10% FBS, 100IU interleukin 2 (IL2) (Peprotech, USA), 10ng/ml IL7 (Peprotech, USA). and IL15 (PeproTech, USA). Then, 2×10^5^ amplified T cells along with αPD-1-(iRGD)_2_ or αPD-1(1, 5, 10, 25, 50μg/ml) along or with free iRGD(100μg/ml) were co-cultured with 2×10^4^ CFSE labeled HGC27 cells for 24 hours. In competitive inhibition assay, the concentration of αPD-1-(iRGD)_2_ or αPD-1 was 10μg/ml, free iRGD was 100μg/ml, αNRP1 was 15 μg/ml, extra αPD-1 was 50 μg/ml. To induce exhaustion of T cells, amplified T cells were cultured in Dynabeads™ Human T-Expander CD3/CD28(Gibco), 500IU interleukin 2 (IL2) (Peprotech, USA), 10ng/ml IL7 (Peprotech, USA). and IL15 (PeproTech, USA). After the incubation, 100ng/ml Propidium Iodide (PI) were added into cultural media. Tumor cell cytotoxicity was measured using flow cytometry. CFSE and PI double positive cells were considered to be lysed tumor cells. The percentage of cytotoxic activity was calculated with the following equation:

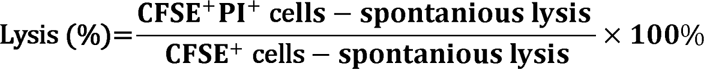

### T cell activation assay

2×105 T cells or OT-I cells were cocultured with 2×10^4^ HGC27 cells or B16-OVA for 24h in 96 well ultra-low adsorption culture plate. The concentration of αPD-1-(iRGD)_2_ or αPD-1 was 10μg/ml, free iRGD was 100μg/ml, αNRP1 was 15 μg/ml, extra αPD-1 was 50 μg/ml. After the incubation, the plate was centrifuged and supernatant was discarded. 100 μL PBS was added into every well to form single cell suspensions. The staining and flowcytometry process were described in “Flow cytometry analysis” part.

### Cell conjugates formation assay

Jurkat-PD-1 cells or OT-I cells were labeled with 2μM CFSE (Abcam, Cambridge, UK). HGC27 cells were labeled with Dye eFluor™ 670 (Thermo Fisher Scientific). In fluorescence imaging, PBMC were labeled with CellTracker™ Deep Red Dye (Invitrogen). HGC27 cells were labeled with CFSE. After washed 3 times with FACS buffer (DPBS containing 2% FBS), 2×10^5^ Jurkat-PD-1 cells or OT-I cells or PBMC were cocultured with 2×10^4^ HGC27 cells. αPD-1-(iRGD)_2_ (10μg/ml) or αPD-1(10μg/ml) along or αPD-1(10μg/ml) with free iRGD(100ug/ml) was added into cultural media. After 1 hour of incubation at 37[, flow cytometry analysis was performed directly using BD Accuri C6 PLUS Flow Cytometry (BD Biosciences). All raw data were analyzed using FlowJo software (10.4, Tree Star). For fluorescence imaging, 1×10^6^ PBMC were cocultured with 1×10^6^ HGC27 cells with 10 μg/ml αPD-1-(iRGD)2 or αPD-1 or indicated agents. If added, the concentration of free iRGD was 100μg/ml, αNRP1 was 15 μg/ml, extra αPD-1 was 50 μg/ml. After 12h, fluorescence figures were taken with ZEN 710 confocal microscope (Zeiss, Jena, Germany).

### Construction and treatment of mouse tumor models

For *in vivo* tumor suppression assay, 1 × 10^6^ MFC or 1 × 10^5^ B16F10 cells were subcutaneously inoculated into 6–8-week-old sex-matched 615-line mice or C57BL/6 mice (n=6 per group). All mice were checked every day. Length of the long axis (a) and vertical axis (b) of tumor nodule were measured with calipers. Tumor volume was calculated via the following formula:

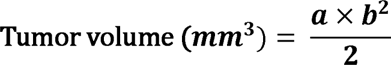

Endpoints of animal experiments and timepoints of tissue collection (tumor and lymph nodes) were shown in figure annotations. In order to obtain single cell suspensions, tumor nodules were shredded and digested with 1mg/ml collagenase IV (Sigma-Aldrich) in serum free RMPI-1640 for 2h at 37[. After the incubation, cells were filtered via 40 μm nylon cell strainers (Biosharp), and washed twice with PBS. Then red blood cell lysis buffer (Biosharp) was applied for erythrocyte clearance. After tumor inoculation, mice were randomized to treatment groups. In tumor suppression assays, mice were treated intraperitoneally (i.p.) with αPD-1-(iRGD)_2_ (5mg/kg), αPD-1 (5mg/kg) along or with free iRGD (1.25μg), free iRGD peptide (1.25μg), 4 times over 12 days.

### Histology analysis

Mouse main organ tissues (heart, liver, spleen, lung, kidney) were prepared as 5 μm formalin-fixed, paraffin-embedded (FFPE) sections. Hematoxylin-eosin staining was done in mouse organ tissue sections, and treatment toxicity was analyzed.

### Flow cytometry analysis

As previously described, single cell suspensions were prepared from tumor tissues and draining lymph nodes. For cell-surface markers detection, single-cell suspensions were stained with antibodies specific to markers including CD3, CD4, CD8, CD25, CD69, PD-1, CD51, NRP-1 for 30 minutes at 4[. For intercellular markers, single cell suspensions were treated with fixation/Permeabilization Solution Kit (BD Biosciences) and/or Leukocyte Activation Cocktail with BD GolgiPlug™ (2μl per 10^6^ cells, BD Biosciences) before the incubating with antibodies targeting IFN-γ and GZMB. Stained cells were examined using BD Accuri C6 PLUS Flow Cytometry (BD Biosciences). All raw data were analyzed using FlowJo software (10.4, Tree Star). The following antibodies were used: anti-mouse CD45 (FITC, #157607), anti-mouse CD3 (FITC, #100203l; PE, #124609), anti-mouse CD4 (PE, #100407; APC, #100516), anti-mouse CD8 (APC, #100711; Percp-Cy5.5, #100732; Alexa Fluor® 700, #100730), anti-mouse CD279 (PD-1) (PE, #135205), anti-CD51(APC, #920011), anti-NRP1(Percp-Cy5.5, #145208), anti-CD25(Percp-Cy5.5, #302626), anti-CD69(APC, #310910), anti-mouse CD25(PE, #12-0251-83), anti-mouse CD69(FITC, #104506) from Biolegend, anti-human IgG(SureLight® PE, ab131612) from Abcam. For intracellular marker expression, single-cell suspensions were stained with anti-mouse IFNγ (APC, Biolegend, #505809; Percp-Cy5.5, #505822), anti-mouse GZMB (AF647, Biolegend, #515406), anti-IFNγ (BV570, #502534), anti-GZMB (FITC, #372206).

### Cell sorting

Single cell suspensions were acquired as previously described and purified by a 67% Percoll gradient (800g at 20 °C for 20 min) to enrich lymphocytes. Then, cell suspensions were stained with Zombie Aqu^TM^ Fixable Viability Kit (#423101, Biolegend) and anti-CD45 FITC (#157607, Biolegend) for 30 minutes at 4[. CD45^+^ Cell sorting was performed using the FACS Aria II (BD Biosciences) system.

### Cell barcoding, multiplexing and scRNA-seq

Generally, one sample was barcoded with ClickTags by the following procedure: for each ClickTag preparation, 0.5 μl of NHS-TCO (1 mM in DMSO) and 20 μl of 25 μM Tz-oligo was thoroughly mixed and immediately pipetted into the sample. After 15 minutes of cell barcoding in the dark at room temperature on a rotating platform, the process was quenched by 10μl of quenching buffer (300 μM alkyne-Tz in FBS) for 5 minutes(47). Then, cells were washed three times with DPBS and detected cell number and viability, before pooled and loaded to a microwell chip targeting 15,000 cells on Singleron Matrix (GEXSCOPE Single Cell RNA-seq Kit, Singleron Biotechnologies, Nanjing, China). Libraries of seRNA-seq were prepared according to manufacturer’s instructions (Singleron Biotechnologies, Nanjing, China). After amplification, cDNA and ClickTags were respectively separated by SPRI size-selection with 0.6× and 1.4× SPRI. ClickTag libraries were subsequently quantified (Qubit, Invitrogen) and amplified using primer SGR-beads-1/SGR-tag-1 and indexed by additional PCR with primer SGR-beads-2/SGR-tag-2. Final ClickTag libraries and transcriptome libraries were analyzed on a BioAnalyzer high-sensitivity DNA kit (Agilent) and sequenced on Illumina NovaSeq 6000. Specially, two-dimensional barcoding was enabled by mixing two different Tz-oligos (10 μL, 50 μM for each ClickTag).

### Analysis of scRNA-seq data

Samples were demultiplexed and aligned using celescope to obtain a raw read count begin of barcodes corresponding to cells and features corresponding to detected genes. Read count matrices were processed, analyzed, and visualized in R (R Core Team, 2013) using Seurat v.4.

## Data availability statement

All of the data supporting the findings of this research are included in the article and the supplementary materials.

## Statistics

GraphPad Prism 7 software were used for statistical analysis of data and graphic representations. For comparisons between two groups, unpaired Student’s t-test, log-rank test or Wilcoxon rank sum test were used as indicated. For gene enrichment analysis, hypergeometric distribution test was used as indicated. The error bars of data were presented as the means ± SEM or means ± SD as indicated. The p value of less than 0.05 was considered to be statistically significant. ns, not significant; *P < 0.05, **P < 0.01, ***P < 0.001 and ****P < 0.0001.

## Study approval

All animal experiments were approved by the Institutional Animal Care and Use Committee of Drum Tower Hospital (approval number: 2020AE01064).

## Author contributions

Conceptualization: JW, JPL, YFP

Methodology: YFP, QX, ZTS, TS

Investigation: YFP, TS, HBW

Visualization: YFP, QX, XRS

Funding acquisition: JW, JPL

Project administration: YFP, QX, ZTS, XYY

Supervision: JW, JPL, YY

Writing – original draft: YFP, QX, ZTS

Writing – review & editing: JW, JPL

The method used to assign the authorship order among the three co-first authors was alphabetic order of last name.

## Acknowledgments

This work was funded by grants from National Natural Science Foundation of China (82073382), the Fundamental Research Funds for the Central Universities (0214-14380506). The funding sources had no role in the study design, data collection, data analysis, data interpretation, or writing of this study.

Thanks are due to Lixia Yu for instruction of in vitro experiments, Fanyan Meng for assistance with the confocal fluorescence imaging and Yang Yang for cooperation with LC-MS analysis.

