## Supplementary Figure1-11 for "Anti-PD-1-iRGD Peptide Conjugate Boosts Antitumor Efficacy via Engagement Augmentation and Penetration Enhancement of T cells"

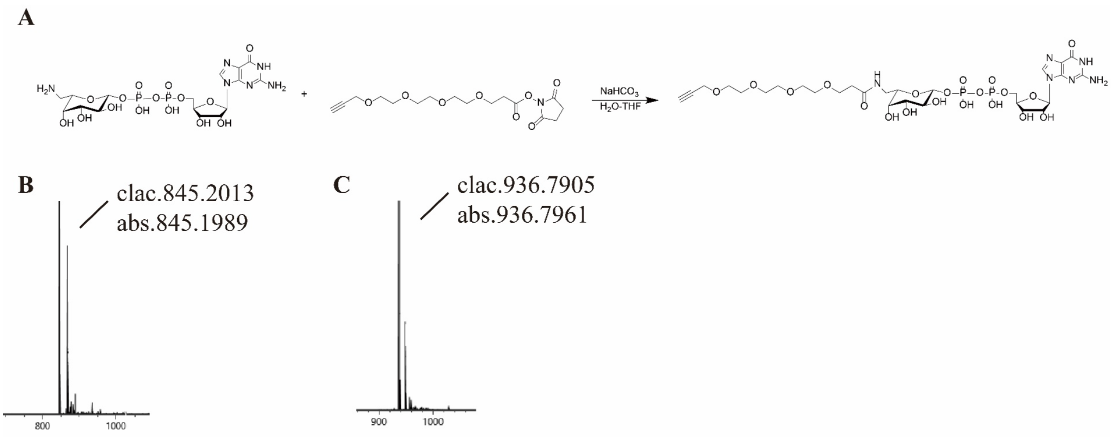


**SI Figure1. Synthesis and characteration of core substrate**

**(A)** Scheme of the synthesis of GDP-FAmP4Prop. **(B)** ESI-MS characterization of GDP-FAmP4Prop. **(C)** ESI-MS characterization of GDP-Fucose-iRGD.

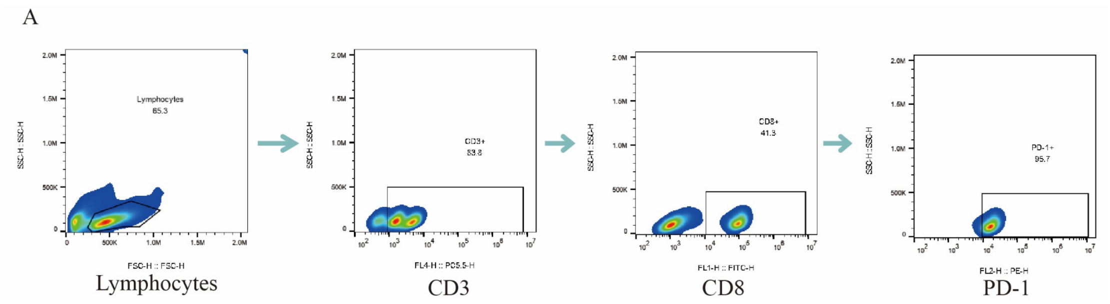


**SI Figure2. Identification of PBMC**

**(A)** Flow cytometry chart of *HLA-A*2402*$^{+}$ PBMC stimulated with CD3/CD28 beads along with IL2 300IU, IL7 10ng/ml, IL15 10ng/ml for a week.

A

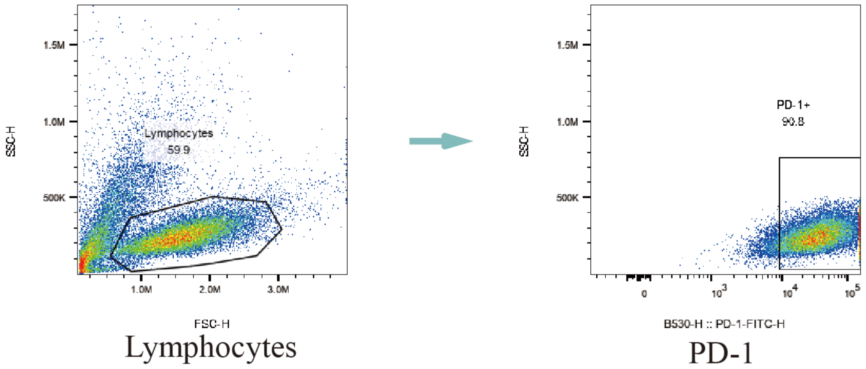


**SI Figure3. PD-1 expression of OT-I**

**(A)** Flow cytometry chart of PD-1 expression on OT-I cells stimulated with IL2 500IU for a week.

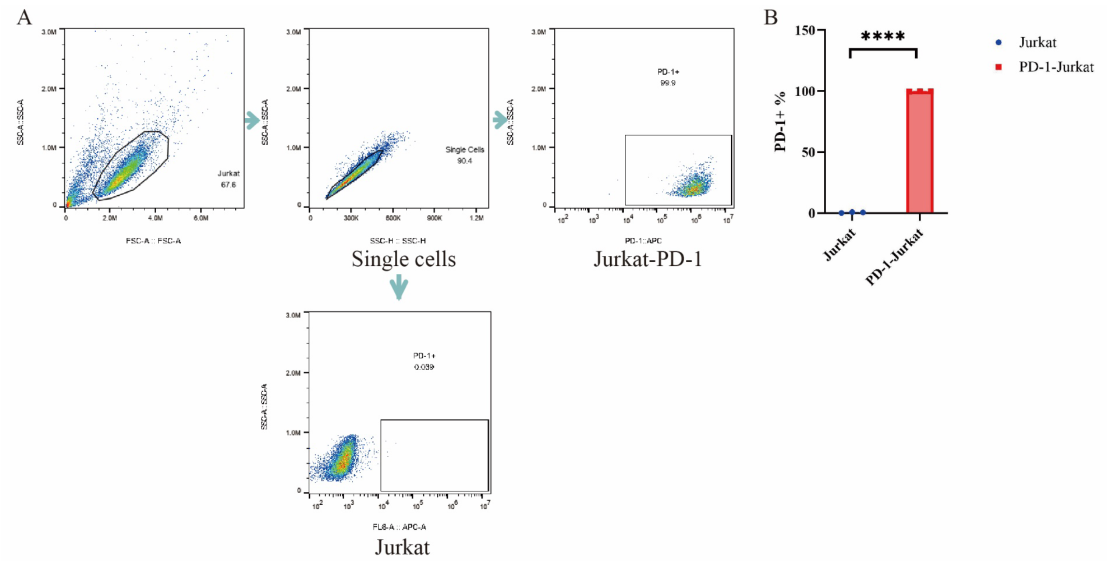


**SI Figure4. PD-1 expression of Jurkat-PD-1**

**(A)** Flow cytometry chart of PD-1 expression on Jurkat and Jurkat-PD-1 cells. **(A)** Histogram of PD-1 expression on Jurkat and Jurkat-PD-1 cells. Data represent mean ± s.e.m.; n = 3. Student's t test. n.s, not significant; *p < 0.5; **p < 0.01; ***p < 0.001; ****p < 0.0001.

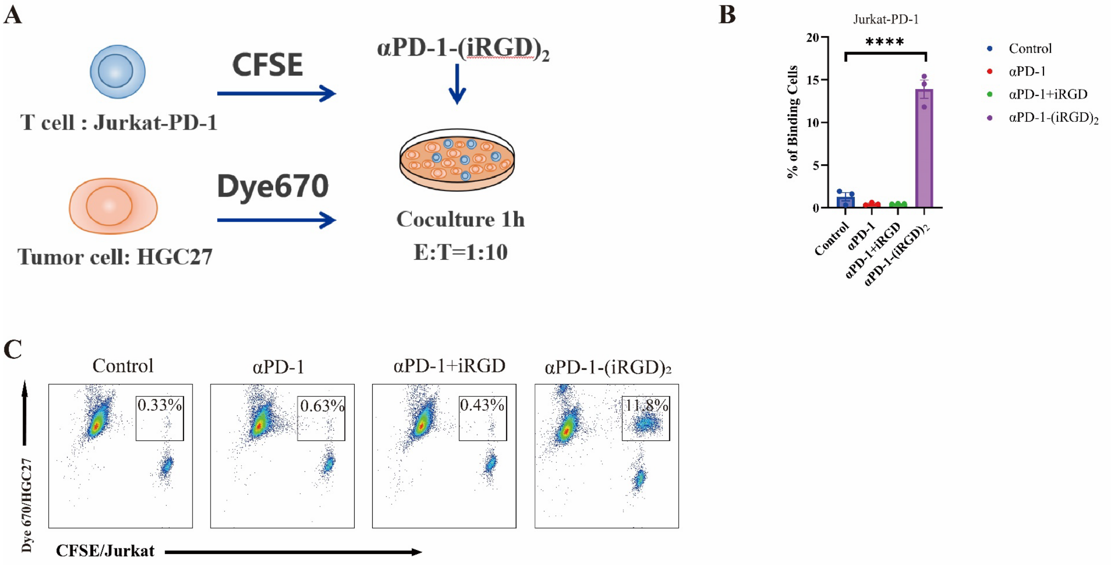


**SI Figure5. αPD-1-(iRGD)$_2$ engages Jurkat-PD-1 and tumor cells**

**(A)** Scheme of the conjugate formation assay of Jurkat-PD-1 and HGC27. **(B)** Percentage of CFSE and Dye670 both positive cells in HGC27 cells incubated with Jurkat-PD-1 cells at an E: T of 1:10. **(C)** Representative flow cytometry results of **(A)**. Data represent mean ± s.e.m.; n = 3. Student's t test. n.s, not significant; *p < 0.5; **p < 0.01; ***p < 0.001; ****p < 0.0001.

**A**

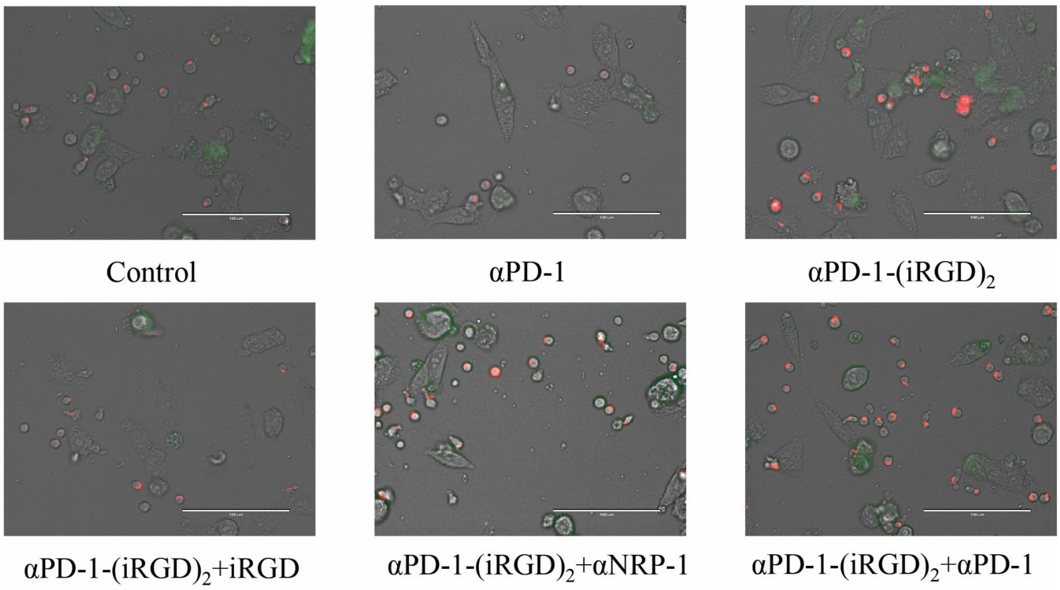


**SI Figure6. αPD-1-(iRGD)$_2$ engages Jurkat-PD-1 and tumor cells**

**(A)** Fluorescence image of 1*10$^6$ CellTracker™ Deep Red Dye-labeled PBMC and CFSE-labeled HGC27 with 10 μg/ml αPD-1-(iRGD)$_2$ or αPD-1 or indicated agents. If added, the concentration of free iRGD was 100μg/ml, αNRP1 was 15 μg/ml, extra αPD-1 was 50 μg/ml.

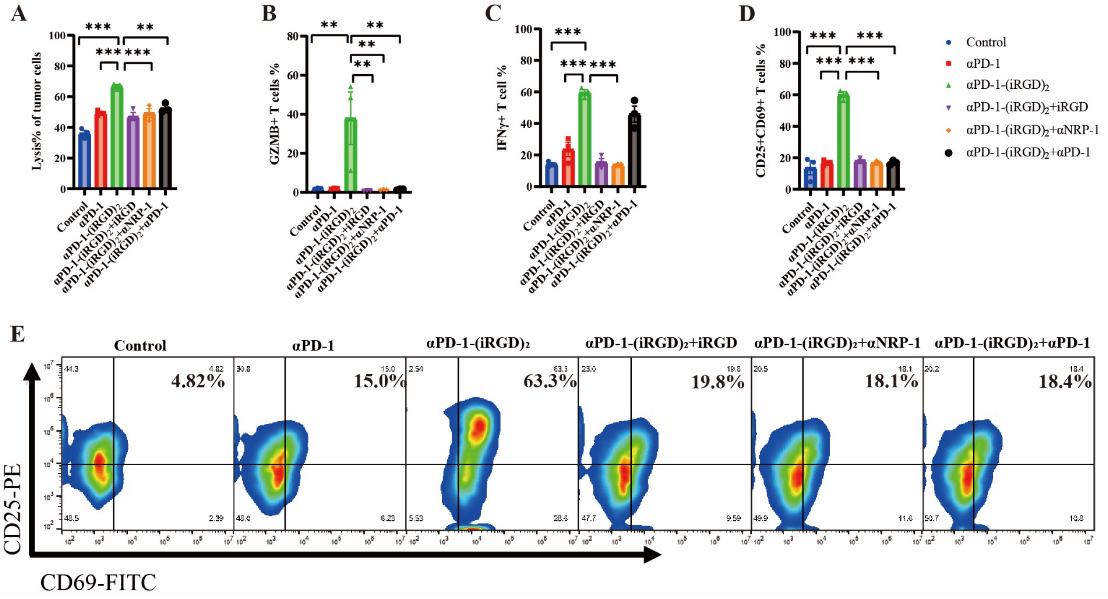


**SI Figure7. αPD-1-(iRGD)2 promoted OT-I cell activation and cytotoxicity**

**(A)**Lysis % of B16-OVA cells coculturing with OT-I cells at an E: T rate of 10:1. Concentration of αPD-1, αPD-1-(iRGD)$_2$ was 10μg/ml, iRGD was 100μg/ml, αNRP1 was 15μg/ml. Specially in αPD-1-(iRGD)$_2$+αPD-1 group, the concentration of αPD-1 was 50μg/ml. **(B)** Expression of GZMB in OT-I cells under coculture conditions in (A). **(C)** Expression of IFNγ in OT-I cells under coculture conditions in (A). **(D)** Expression of CD25 and CD69 in OT-I cells under coculture conditions in (A). **(E)** Representative flow cytometry results of (D)**.** Data represent mean ± s.e.m.; n = 3. Student's t test. n.s, not significant; *P < 0.5; **P < 0.01; ***P < 0.001.

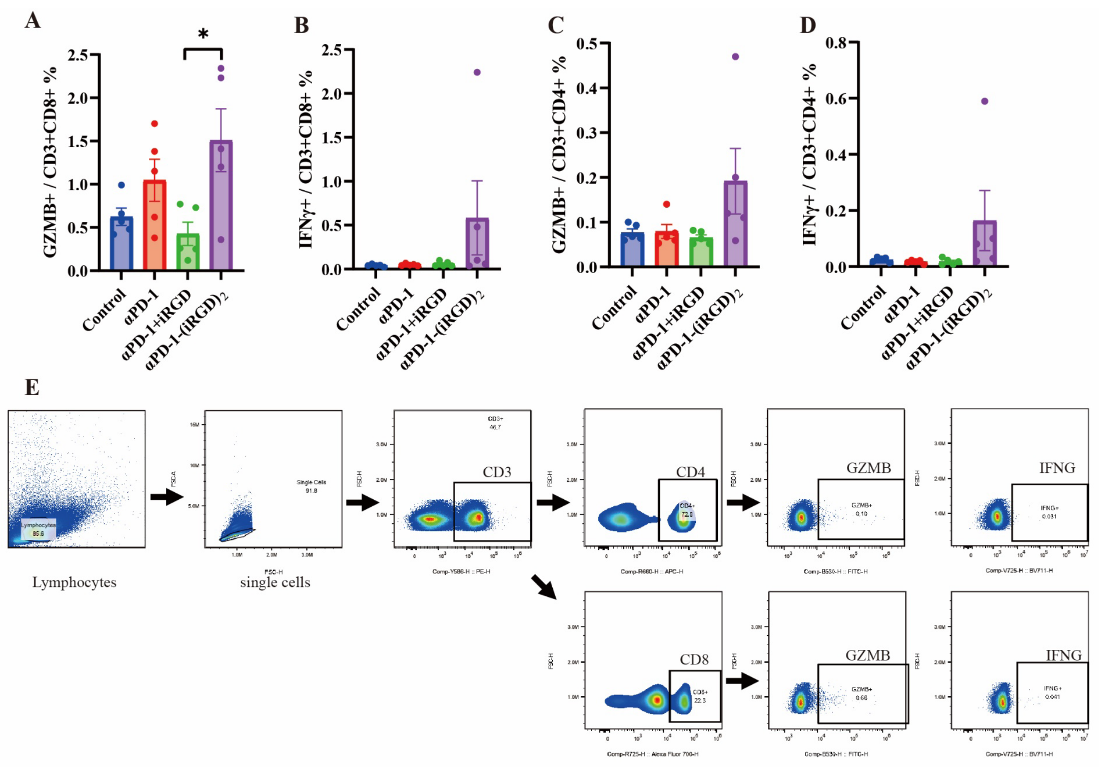


**SI Figure8. αPD-1-(iRGD)₂ activates $CD4^{+}$ and $CD8^{+}$ T cells in tumor draining lymph node**

**(A)** Quantification of GZMB expression on $CD8^{+}$ T cells from resected draining lymphnodes at the end point of MFC tumor bearing mice treated with αPD-1-$(iRGD)_2$ or control reagents. **(B)** Quantification of IFNγ expression on $CD8^{+}$ T cells. **(C)** Quantification of GZMB expression on $CD4^{+}$ T cells. **(D)** Quantification of IFNγ expression on $CD4^{+}$ T cells. **(E)** Gating strategy of flow cytometry. Data represent mean ± s.e.m.; n = 5. One-way ANOVA; n.s, not significant; *P < 0.5; **P < 0.01; ***P < 0.001; ****P < 0.0001.

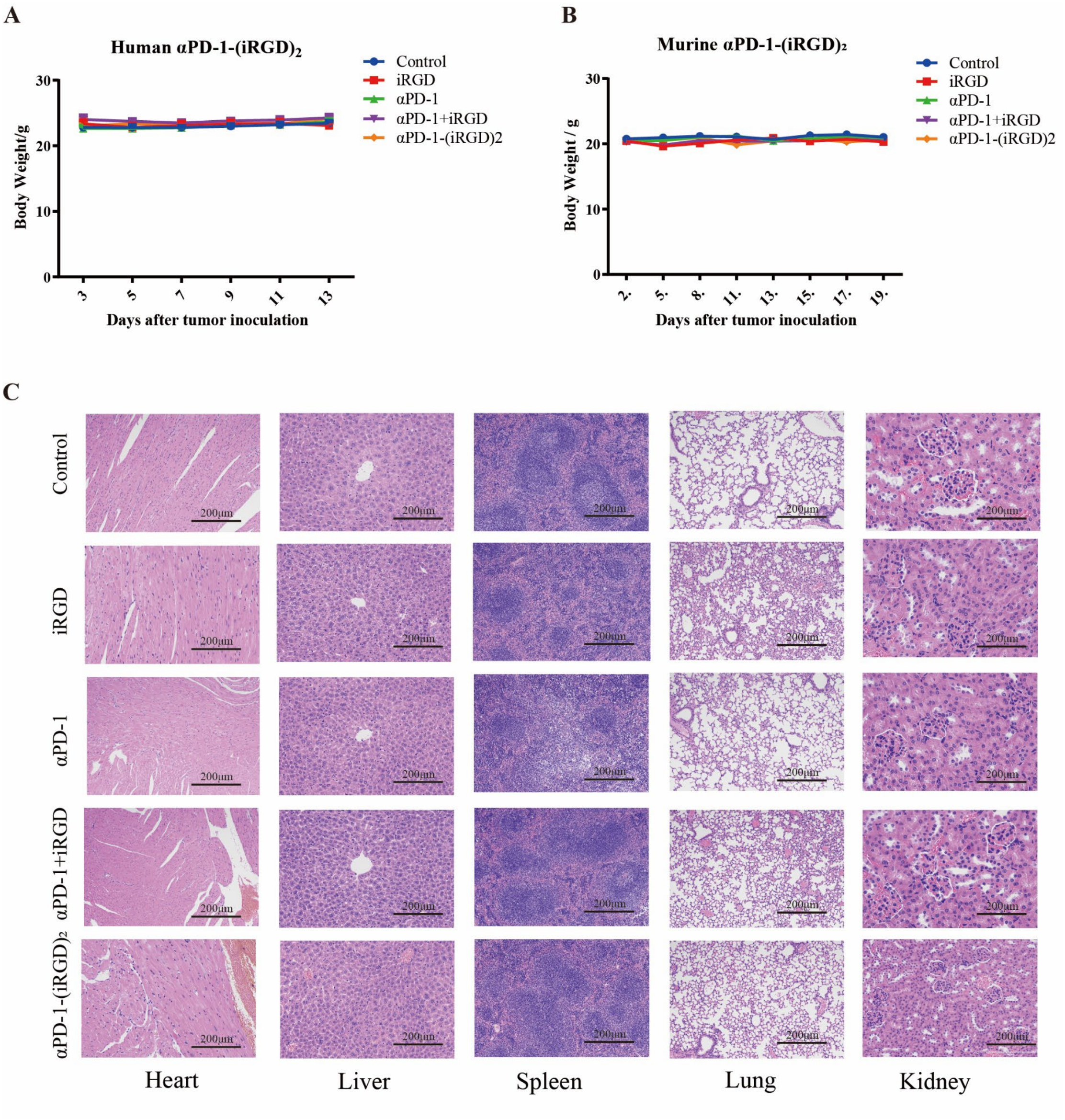


**SI Figure9. Biosafety analysis of αPD-1-(iRGD)$_2$**

**(A)** Body weight plot of MFC tumor bearing mice after treated with αPD-1-(iRGD)$_2$. **(B)** Body weight plot of MFC tumor bearing mice after treated with murine αPD-1-(iRGD)$_2$. **(C)** H&E staining of major organs in MFC subcutaneous mouse model after the treatment of αPD-1-(iRGD)$_2$.

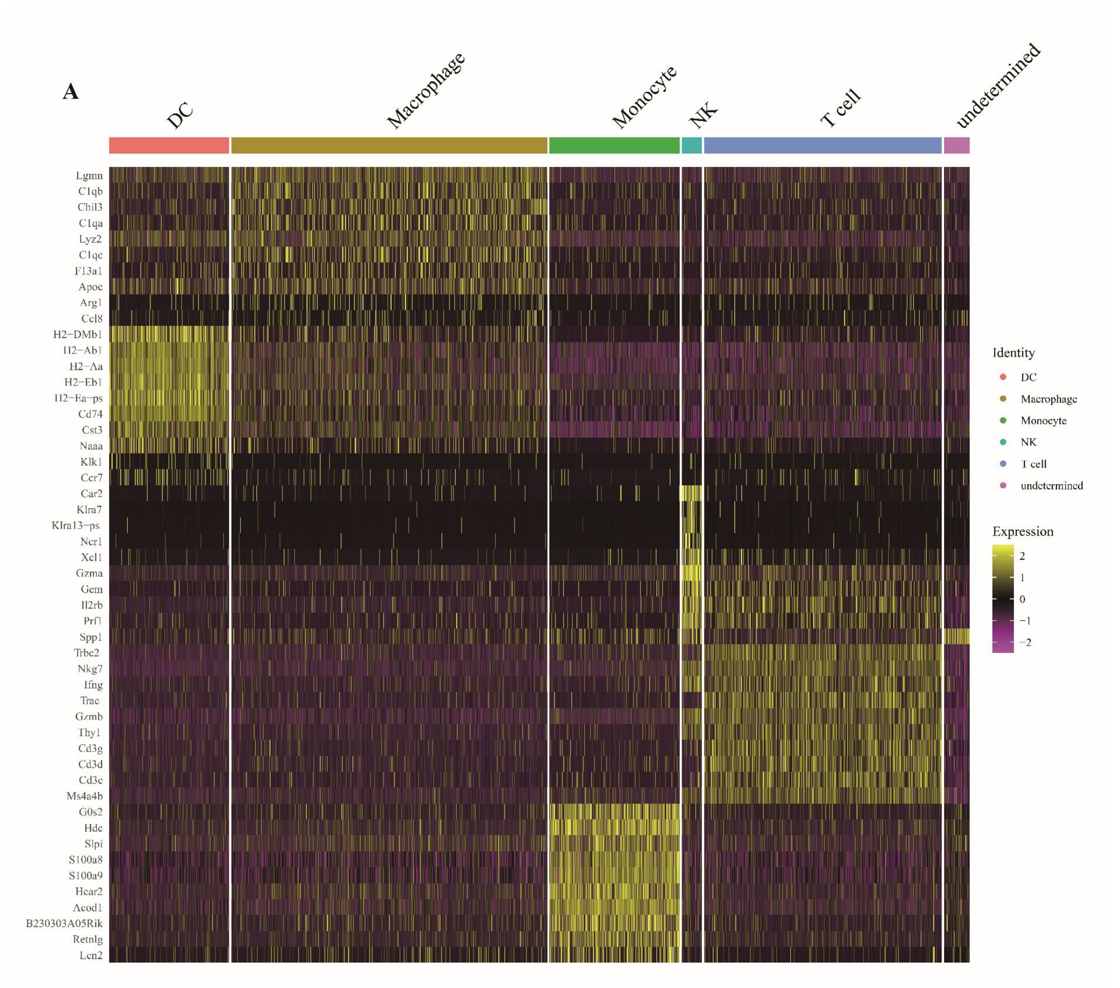


**SI Figure10. Clustering markers of two-dimensional (2D) UMAP visualization of CD45⁺ tumor infiltrating immune cells**

CD45$^+$ cells were collected from MFC tumor bearing mice treated with αPD-1-(iRGD)$_2$.

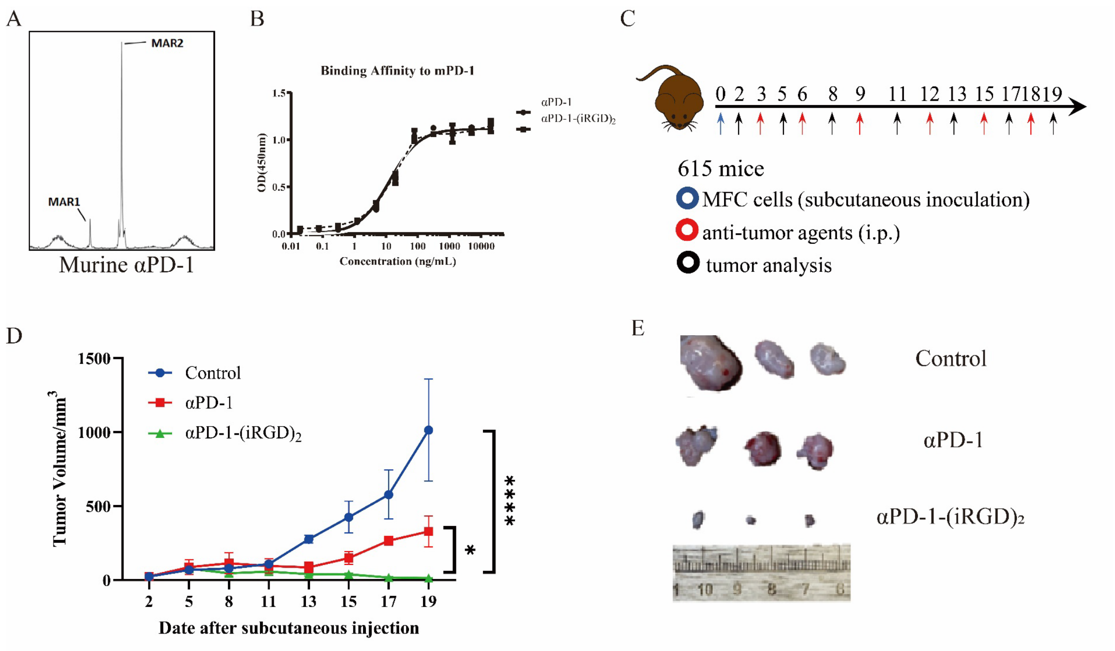


**SI Figure11. Synthesis and antitumor efficacy of murine αPD-1-(iRGD)$_2$**

**(A)** ESI-MS characterization of murine αPD-1-(iRGD)$_2$. **(B)** Binding affinity of murine αPD-1-(iRGD)$_2$ and unmodified antibody towards murine PD-1 protein by ELISA. **(C)** Schematics of the treatment regimen in MFC homograft models are shown. Briefly, mice bearing tumor burdens were treated with $1 \times 10^6$ MFC cell and injected intraperitoneally with PBS (100ul control), αPD-1(10mg/kg) or αPD-1-(iRGD)$_2$ (10 mg/kg) every three days. **(D)** Tumor growth profiles of **(C)**. **(E)** Ex vivo imaging of tumors at end point. Data represent mean ± s.e.m.; n = 3. Two-way ANOVA, n.s, not significant; *P < 0.5; **P < 0.01; ***P < 0.001; ****P < 0.0001.
